# Advancements in Developing an Automated Breast Density Detection Technique for Breast Cancer Risk Prediction: Synthesizing a Signal-dependent Noise Stochastic Process

**DOI:** 10.1101/2025.11.25.690493

**Authors:** John Heine, Erin Fowler, Matthew Schabath, Kathleen M. Egan

## Abstract

Breast density is an important breast cancer risk factor estimated from mammograms and useful for breast cancer risk prediction. We describe a new formulation built on the backbone of an established automated percentage of breast density detection method. This framework relies on signal-dependent noise (SDN), characterized by a statistical dependence between the variance and mean signal. Variations in this dependency due to different image data representations cause degradation in the algorithm’s performance; the current work addresses this problem by synthesizing a stochastic process conditioned on a given mammogram instead of analyzing the mammogram directly. Image data used in the analysis was derived from three breast cancer case-control studies employing different mammographic technologies including full field digital mammography (both raw and clinical images) and digital breast tomosynthesis. The new formulation produced significant odds ratios across all image data representations due to these methodological advancements: (1) synthesis of SDN to an optimal quadratic structure given an arbitrary image; and (2) ensemble averaging over a given image, boosting the signal. We also demonstrate methods to standardize and combine measurements from different technologies using a probability density transformation technique. This automated technique can be applied to images from different technologies with minimal adjustment, thereby making it suitable for both research and clinical applications.

## Background

Breast density is observable in mammograms with or without automation and is a widely accepted significant breast cancer risk factor [1, 2]. The aim of the present work is to introduce a new formulation founded upon an established automated breast density detection algorithm [3–5] using digital mammograms acquired in multiple data representations. Inclusion of this image measure in models that predict absolute risk over time may be useful for personalized breast screening strategies [6–8], thereby facilitating epidemiologic and clinical research on this important breast cancer risk factor.

In mammograms used for clinical purposes, breast density is loosely defined as radiographically *bright* tissue. More specifically, mammograms are typically modeled as a *two-component* system comprised of adipose and glandular breast tissue (i.e., breast density). For the x-ray energies used in mammography, this two-component model is an appropriate approximation. Nevertheless, dense tissue is a combination of both the at-risk epithelial (ductal and lobular) and stromal (supporting or connective) tissues [9, 10], which are likely inseparable in x-ray mammograms due to their similar x-ray attenuation coefficient values relative to the respective x-ray energies.

The user-assisted measure of breast density, referred to as Cumulus (University of Toronto, Toronto Canada) [11–13], is considered the benchmark for comparison with other measures [14]. This technique computes the percentage of dense tissue within the breast area. In Cumulus, the operator performs the following tasks based on brightness and contrast adjustments: (1) segments the breast area from the background; and (2) sets a threshold, labeling everything above as dense and below as adipose tissue. This results in a binary labeled image within the breast area and yields an estimate of the proportion of the tissue equivalent to the percentage of breast density (PBD). This user-assisted measurement is resource-intensive, and both segmentation and labeling thresholds are subject to individual user preference, and their assignment prone to random intra-user variation. In contrast, automation should remove subjectivity, and provide a more reproducible measurement, as well as more timely and less labor-intensive breast density estimate in clinical and research applications

The history and merits of quantitative breast density measurements are extensive [15–17]. Image-analytics have evolved to include automated features that may not be precisely visually observable in mammograms. For example, these measures could be generated with filtering or other techniques that summarize spatial relationships [18, 19] as surrogates for breast density or possibly standalone independent predictors of risk [20]. Such measures are often collectively referred to as texture features. More recently, texture-type risk prediction features and models have been generated with more advanced machine learning and artificial intelligence (AI) techniques [7, 21–24]. In some AI applications, convolutional neural networks (CNNs) leverage texture in a unique manner. These networks can be interpreted as hierarchically learned composite filters derived through multiple processing stages [25, 26] that facilitate texture extraction. Although these AI applications show much potential and improved risk prediction capabilities, these latent feature use for decision purposes are problematic because their description is not readily available, hindering clinical adoption [27]. Ongoing research aims to address this limitation by developing methods to interpret features learned by CNN architectures [26].

A major obstacle in developing a universally applicable automated method for breast density estimation when incorporating both current and historical data is the evolution of digital mammography over time: from digitized screen-film images, advancing to full field digital mammography (FFDM), and then to the current digital breast tomosynthesis (DBT). Each technology is associated with unique image data representations and accompanied by an array of distinguishing technical features: different film-digitizers; enhancements for clinical purposes; mammograms acquired from different FFDM detector designs; and addition of a quasi-third dimension with units that produce varied synthetic two-dimensional images, differing both within and across manufacturers. When building a universally applicable risk prediction model that includes breast density, these potential variants must be accommodated. A prime research example of the evolution in technology and ensuing challenges comes from the Nurses’ Health Study (NHS). Originating in the mid-1970s [28, 29] the NHS spans the entire era of mammographic screening and the adoption of new mammography advancements over 40-plus years [30, 31]. In research based on the NHS, inclusion of mammography data from the early years of the cohort necessitated the physical collection and then digitization of film images for thousands of mammograms as a prelude to image analytics [32].

The Breast Imaging and Reporting Data System (BI-RADS) ordinal breast density assessment method (i.e., the breast composition descriptors described in the American College of Radiology Atlas 5^th^ Edition, 2013) [33] has also been used for risk prediction (these breast composition descriptors were also described in earlier editions of this atlas). This mammography reporting technique trades measurement resolution (i.e., a continuous scale) and, in theory enhanced risk prediction potential, for contrast (i.e., a four-state ordinal scale) and ease of use in the clinical workflow. Although initially not developed for risk prediction purposes, it is commonly used for this application [34–36]. The transition of an automated technique to clinical applications from research should require minimal adjustments and have the capability to accommodate a range of image types currently used in clinical practice as well as those anticipated in the future. As an alternative, a viable automated measure should be equally useful in research and clinical environments when compared to the BI-RADS measure.

Algorithmic advancements investigated in this report are intended to produce a more stable PBD measure with the potential for universal application. This measurement operates by analyzing signal-dependent noise, distinguished by a statistical dependence between the variance and mean signal. In past analyses, this method’s performance varied depending on the image data representation (i.e., image format). In radiographic imaging systems, such as mammography, quantum noise (i.e., Poisson noise) is one source of variation [37, 38] that is signal-dependent. Such noise produces a grainy appearance especially noticeable in underexposed image areas. At small spatial scales, noise variation is approximately signal-dependent (i.e., quantum noise) in FFDM units [3, 39]. We have noted that the statistical structure of the noise variance affected the method’s performance [3]. For example, under ideal local conditions, the variance depends on the tissue type: a larger variance for dense tissue and a considerable smaller variance for adipose tissue. In practice, these relationships did not offer sufficient contrast between tissue types in some image formats. In these situations, ad-hoc adjustments were implemented with no attempt to arrive at an optimal algorithm correction [3, 4].

Although the advanced algorithmic formulation described here is built on the foundation of an established methodology, it offers a new breast density detection algorithm with additional operating procedures. Our modified formulation no longer relies on a given mammogram’s intrinsic noise structure. Instead, the detection is based on a synthesized stochastic process that provides a residual image that is analyzed, conditioned on the mammogram. This yields a signal-dependent noise structure that enhances the algorithm’s performance, regardless of a given mammogram’s intrinsic, or natural, noise characteristics. This synthesis follows from a signal-dependent stochastic model that modulates a given mammogram with random noise that supports ensemble averaging, boosting detection performance. Supporting derivations in the Methods Section illustrate the expected variance contrast gain between tissue types by synthesizing the signal-dependent noise in accord with this stochastic model. This synthesis approach added two additional free parameters to the algorithm. As demonstrated, these algorithmic advancements should produce a more stable PBD estimation regardless of image format differences as described below.

In the context of the above-described new formulation, we evaluated the technique’s risk prediction capabilities over varied image formats. The research uses three breast cancer matched case-control studies, each study accompanied by images from different mammographic technologies in use at this center between 2007-2022. The primary objectives were to: (1) evaluate how to determine the operating characteristics of the new detection process over a range of image formats; (2) investigate how to process PBD measures that yield consistent results across studies; (3) develop a standardized approach to facilitate aggregation of data derived from different mammography technologies, a frequent goal in epidemiologic research, and finally; (4) provide a more theoretical framework for the detection formulation and advancements within the context of synthesizing a stochastic process for a given image with a specified SDN structure.

## Methods

### Automated breast density detection technique

The automated density detection techniques was established previously [3–5], and its operational details are provided for context when describing the new formulation; here, the original operational mechanics are integrated into the new formulation. The method produces an output like Cumulus and is referred to as PD_a_ (i.e., automated PBD); this abbreviation is used as reference to both the technique and the measure it produces. It operates by analyzing signal-dependent noise (SDN). In images formed by x-ray fluence, the signal captured by an ideal linear detector would be represented by Poisson statistics at the pixel level (i.e., neglecting other noise type contributions). In this idealized situation, the expected mean scales statistically with the variance when considering ensemble expectations; although, this may not be the most suitable noise structure for the detection process. This ideal Poisson construct approximately holds locally in practice with FFDM images in some formats but not universally. In this context and to a good approximation, the noise variance depends linearly with the mean signal over small spatial scales in both direct and indirect x-ray detection FFDM technologies for raw images used in this study [3, 39]. It is important to recognize, ensemble expectation quantities require acquiring multiple realizations of a given breast (or image) in succession; this implies acquiring multiple mammograms of the same participant without repositioning, changing the x-ray tube orientation, changing the x-ray beam, or changing the breast compression, which is essentially impossible. The modified synthetic advancement overcomes this limitation.

For the density detection, a high-pass filter is applied to a given mammogram to separate it into two image components: the residual high frequency fluctuations, as an approximation for the SDN component, and the lower resolution structure (the signal), which is not used explicitly. The density detection is performed in the residual image. The high-pass filter type has changed since the technique’s inception for algorithm processing time reductions, while serving the same purpose. Because there is only one mammogram per participant (i.e., per breast considering its laterality and orientation) for a given time point normally, ensemble expectations are approximated with local spatial averages taken over relatively small image regions. This approximation is valid because mammograms have long range positive spatial correlation (discussed below). The high frequency residual image is scanned with n×n pixel search window (n∼2-4 pixels). At the local level, using these pixels in an expectation calculation approximates acquiring n^2^ consecutive mammograms of a given participant without repositioning and taking the expectation of a given quantity using n^2^ realizations. In each window, the local variance is calculated and compared with the global adipose reference variance using a chi-square test (two-sided) specified with a preset significance level (defining detection parameter 1). The reference variance is estimated the first time using all pixels in the filtered image within the breast area (see below for an important qualification) in the calculation. To capture the rarer very dense breast event, the detection is repeated after the first stage is completed by setting a second significance level (defining the second detection parameter) and updating the adipose reference variance. The updated and refined reference variance is calculated in the high-pass filtered image using pixels declared non-dense in the first detection stage. This second detection stage produces the PBD binary output image. The second detection stage is required especially when an upper-density image is under analysis. In this situation, the first adipose reference estimation will be less than optimal. In this new approach, the window is *centered* on a given pixel, and the decision is made approximately at the central pixel location (i.e., corresponding to statistical decision at each pixel within a given mammogram). In either stage, when the local variance deviates significantly from the reference, the central pixel is labeled as dense or otherwise it is labeled as adipose. The window is shifted in pixel increments across the filtered mammogram’s breast area applying the same statistical testing method. When operating on images in a different format, detection parameters were adjusted with training data, where case-control data was preferred but not necessary.

For context, PD_a_ is defined as detection technique for the reasons outlined here. In classical (Neyman–Pearson) detection theory, a true detection event (ideally dense tissue) is defined relative to a fixed false-alarm probability, which in this setting corresponds to labeling adipose tissue as dense. In our variant of this approach, a mammogram is transformed into a form suitable for applying a standard statistical test using the ratio of two variances. This serves as a practical surrogate for the Neyman-Pearson likelihood ratio test because the statistical characteristics for dense tissue are not explicitly known [40]. The detection is dependent upon estimating the adipose reference variance used in the ratio test. Supporting work for the two-stage reference variance determination comes from a study that showed a 50% dense breast by volume is rare but exists [41]. In a similar sense, recent work demonstrated how the volumetric breast density structure from DBT projects onto a plane to form the PBD measure [42]; these two studies taken together show why the first adipose reference variance calculation is probably an acceptable estimation for many images that requires fine tuning.

Over the course of our research in developing PD_a_, a simple advancment was overlooked that eliminates two algorithm *restrictions*: (1) dependence on the intrinsic SDN structure of a given image format or image; and (2) approximating the SDN component with high-pass filtering. Because the SDN structure was not always conducive to the algorithm’s performance, image transformations were explored on a somewhat ad hoc basis [3, 4]. Yet, the optimal SDN structure was not understood, and the algorithm could not produce significant findings with some image formats. In this new formulation, the SDN residual image is synthesized within statistical limits to a predictable structure, eliminating the need for high-pass filtering altogether. In pursuit of our goals, the methodology was never evaluated over the full range of detection parameters in the context provided in this current study. An unstated innovation is that the SDN synthesis enables the application of the appropriate statistical test.

### Modified residual detection input image

The main breakthrough follows from changing the detection input image derived from an adaptable stochastic framework. An image corrupted by SDN can be modeled as follows

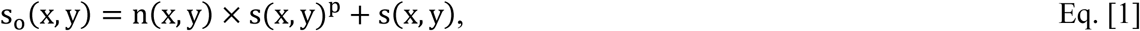

where s_o_(x, y) is the observed noise-corrupted image, s(x,y) is the ideal noiseless image, n(x, y) is random zero mean noise with unit variance in our modeling, 0 < p ≤ 1, and (x, y) are spatial pixel coordinates. Additive random noise has been neglected in Eq. [1]. Variants of this stochastic model have been used in a wide field of investigations as reviewed elsewhere [43]: (1) images formed with charge-coupled device and complementary metal-oxide semiconductor arrays; (2) ultrasound images; and (3) images constructed from synthetic aperture radar. Taking the expected ensemble averages from Eq.[1] gives the mean and variance, respectively: using E[s_0_(x, y)] = s(x, y), yields E[s_0_(x, y)^2^] - E[s_0_(x, y)]^2^ = s(x, y)^2p^, where E[*·*] is the ensemble expectation operator. For n(x, y), we used a standardized normal distribution (zero mean, unit variance). For example, Poisson noise is modeled when p = ½, gives s(x, y) as the expected ensemble mean and variance. In this context, this formulation is interpreted as a stochastic process rather than a static expression. When p = 1, the mean and noise variance have a quadratic structural relationship. The key point here is recognizing the SDN component of the stochastic process in Eq. [1] is a residual image that could be synthesized if s(x, y) was known.

The original detection approach applied a high-pass filter to a given mammogram to approximate the SDN residual image. When working with a single image, this approximation is justified; for a given realization of the stochastic process in Eq. [1], the power spectrum of the residual SDN image is statistically uniform, analogous to that of white noise, and the two-dimensional high-pass filter used here (i.e., a wavelet filter) captures three quarters of the noise variance when applied to such noise when applied to a single realization. Consequently, this provides an acceptable approximation for the residual when filtering images, such as mammograms that approximately satisfy Eq. [1]. Notably, this filter-variance relationship applies to both traditional two-dimensional white noise images and the SDN residual image in Eq. [1] [43]. Because spatial averaging is still used and in furtherance of the approximation, the power spectrum of mammograms follows an inverse power law [5, 44–46]. This characteristic gives rise to long range positive spatial correlation implying mammograms vary slowly spatially. Accordingly, the high frequency residual image approximation, derived from filtering, often has parasitic pixel values relative to the respective mammogram’s intensity range.

In the new algorithm, the detection input image is synthesized using the SDN component from Eq. [1], in isolation, expressed explicitly as the residual component referenced to above

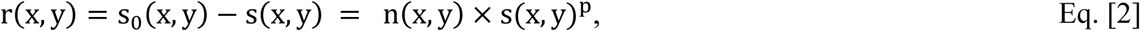

where p > 0 (eliminating the restriction from above), although only two values of p (½ and 1) were investigated. Accordingly, the signal-dependent noise is synthesized using Eq. [2] with a suitable estimation of s(x, y). In this formulation, the residual image is analyzed directly rather than using the filtered image approximation, defining the principal advancement and change in the breast density detection algorithm introduced here. The exponent p is the first additional free parameter in the modified formulation, and its value plays a critical role. The detection is applied by squaring the residual image in Eq. [2] and performing ensemble averaging yielding a chi-square image

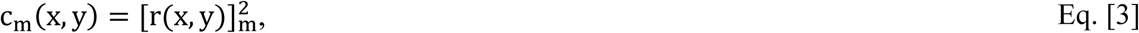

where the integer subscript m > 0 specifies the ensemble size used in the averaging. A given image in this ensemble (specific to one patient) is synthesized using a different realization of the random field n(x, y) conditioned on the the same s(x, y) in Eq. [3]. To approximate s(x, y), a 7×7 boxcar averaging (smoothing) filter is applied to mammograms to suppress high frequency fluctuations. Averaging m ensemble realizations produces the image defined in Eq. [3]. Because E[r(x,y)] = 0, ensemble averaging yields a scaled chi-square variate with m degrees of freedom at each pixel (as qualified in the *Chi-square image statistical characterization* section below), collectively expressed in Eq. [3]. This advancement enables averaging over realizations of the squared residual image and is thereby interpreted as a stochastic process; this is the second fundamental change in the reformulated methodology, adding a second degree of freedom in the advanced algorithm. Because m spans an unlimited number of possibilities, we investigated specific values: (1, 5, 25, 50 and 100). This synthetic construction becomes essential when the detection input image is highly processed or in other situations where Poisson or SDN statistics may not hold. Such situations may arise from heavy processing used in DBT reconstruction, images with a compressed density-adipose intensity scale, or clinical images that have been enhanced with nonlinear techniques.

The structural form of a given mammogram also plays an important role in the detection process; in this context, the chi-square statistic applicability and form are discussed. In theory, starting from Eq. [2] together with Eq. [3] ensures there is a predictable SDN relationship for a given image format at (x,y) Although derived by a smoothing operation, the underlying structural variations in each mammogram are not substantially affected in the noiseless s(x, y) approximation; local structural variations can contribute to the local variance computed within the search window and can perturb the chi-square distribution applicability. However in favor this approximation, the scaled chi-square variance estimate is appropriate when ensemble averaging is followed by spatial averaging because the residual image r(x, y) exhibits negligible linear spatial correlation [43] and mammograms vary slowly spatially (discussed above). When scanning c_m_(x, y) with the search window, the chi-square detection in the new approach uses the average of n×n pixels about position (x, y). The following spatial averaging produces approximately a chi-square variate with (n^2^-1) × m degrees of freedom at (x, y), where the degree of freedom reduction follows from calculating the local variance in the detection stages. The selection of n for a given image format is discussed below. We refer to an image formed by ensemble and spatial averaging of the squared residual image as a *chi-square image*.

### Chi-square image statistical characteristics

In the new formulation, ensemble expectation quantities are calculated from the scaled chi-square distribution to understand their impact on the detection performance; the mean (µ) and standard deviation (σ) derived from Eq. [3] are discussed here. In Eq. [2], we make the substitution d = s(x,y)^p^ for a fixed pixel location and use this substitution in Eq. [3] with m = k corresponding to averaging k independent and identically distributed chi-square variates with one degree of freedom at that (x, y) location. This yields a scaled chi-square variate with k degrees of freedom with expected mean and standard deviation given by µ = d^2^ and 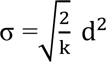. Showing in principle, the uncertainty (i.e., σ) in the ensemble averaged estimator at a given pixel can be made arbitrarily small by increasing the number of realizations; this refers to the random fluctuation about the structural form. The limiting value in µ follows from E[n(x, y)^2^] = 1. Accordingly, in the limit k → ∞, the SDN amplitude becomes parasitic, paradoxically limiting detection performance (as discussed in results). To illustrate the SDN structure of a given chi-square image realization, the local variance is estimated relative to its mean signal. This is achieved by applying a spatial averaging technique that approximates ensemble averaging across all spatial locations, yielding a SDN plot [39]; these plots show the structural SDN from with the variation suppressed to varying degrees due to the averaging process in the respective algorithm. Clinical mammograms are used as comparators by computing their intrinsic SDN curves obtained by the same plot-generation methodology, but the high-pass filtering approximation is employed as in the original formulation.

### Additional preprocessing for raw mammograms

Prior experiments indicated raw Hologic FFDM images required a non-linear transformation to facilitate the density detection [3]. Prior to this investigation, we explored alternative transformations to raw mammograms acquired from different imaging technologies. Raw FFDM images at this center are stored in monochrome 2 format: dense tissue appears dark and adipose tissue appears bright. First, after the boxcar filter application, the intensity scale is reversed by subtracting the (pixel bit-rate -1) from a given image. This reversed scale image is normalized by dividing by its maximum pixel value yielding t(x, y). The ideal noiseless image approximation is then defined as s(x, y) = exp [c × t(x, y)]. The constant, c, is computed by setting the largest pixel value in each s(x, y) image to 16,000, a heuristic choice based on prior experimentation. The same transformation was used for both Hologic and GE raw FFDM images. This transformation is based on prior experimentation and not claimed to be globally optimal.

### Algorithm qualifiers

For algorithm replication purposes, the breast area was eroded radially inward [47] (i.e., treating the breast area as an approximate half hemisphere) by 2% before estimating reference values and performing the detection. This is to mitigate edge effects in each image. Such effects can skew both global and local variance calculations. The full breast area was included in the normalization for PD_a_.

### Residual image variance dependency on p

The image modeling can be related back to both the x-ray image acquisition, when it applies to a physics-based model, and to image processing, when the acquisition derivation is not applicable. Both formulations rely on the expectation values for the residual image in Eq. [2] to illustrate the role of the exponent p.

The first approach is related to an idealized x-ray image acquisition model using Beer’s Law to investigate the role of the exponent p in the equations [1 and 2]. This approach applies strictly to raw intensity images. The transmitted intensity is expressed as

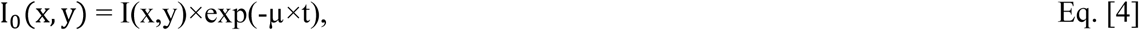

where μ is an effective linear x-ray attenuation coefficient, t is the compressed breast thickness, I(x,y) is the spatially uniform incident x-ray beam, and I_0_(x, y) is the idealized acquired image. Because x-ray transmission follows Poisson statistics, Eq. [4] represents the mean expected transmitted intensity with variance proportional to the mean. For typical Hologic FFDM acquisitions (W/Rh spectrum, 25 - 30 kV), measured representative effective linear x-ray attenuation coefficients are: µ_a_ = 0.45cm^−1^ for adipose tissue and µ_d_ = 0.60cm^−1^ for dense tissue [48]. These values with t = 6cm (approximately, the average compressed breast thickness) are used to illustrate the role of p. In the native raw image intensity domain (non-inverted intensities, where adipose tissue appears bright and dense tissue dark), the relative separation between the two types of tissue scales as

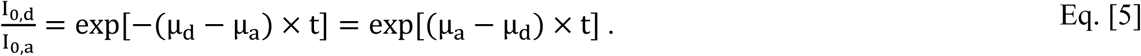

Because our analysis is performed on intensity inverted images, only the magnitude of the separation enters the analysis. Defining Δµ = |µ_𝑎_ − µ_𝑑_| > 0, the relative separation scales as exp (Δµ × t). To compute the variance separation while incorporating p from Eq. [1], the relative variance gain is given by:

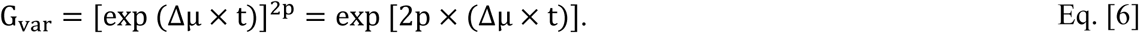

Using the representative values from above gives the relative gain:

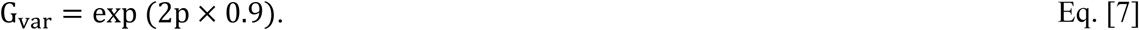

For the values for p investigated in this report: G_var_ ≈ 2.5 ≈ when p = ½ and G_var_ ≈ 6.0 when p = 1 (producing approximately 2.5 relative gain for the quadratic structure). Moreover, using measured effective linear x-ray attenuation coefficients from a GE unit (Mo/Mo spectrum over similar rage of kV), with u_d_ ≈ 0.81, u_a_ ≈ 0.56, and t = 6, yields G_var_ ≈ 4.5 for p = ½ and G_var_ ≈ 20 for p =1[48] (producing approximately 4.5 relative gain for the quadratic structure). Relating back to Eq. [2], this analysis suggests increasing p should produce a substantial amplitude difference between the adipose and glandular tissue variances. In our experiments, this increased variance gain is expected to manifest as large (or significant) ORs in logistic regression modeling.

The second approach applies to analyzing the residual image in settings where Beer’s law does not strictly apply, such as with the synthetic 2D images from DBT, when non-linear processing was applied to the raw data to produce clinical images, or for other settings that may arise; it also serves as an alternate framework when Beer’s law applies. We work with conventional intensity values and assume the baseline pixel values for dense and adipose tissue satisfy s > s . From the residual image, the variances are expressed as 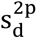 and 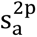, respectively.

The relative gain is therefore:

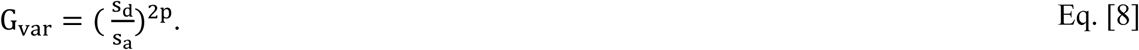

Equivalently in logarithmic form,

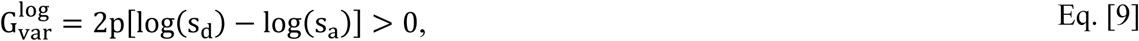

demonstrating the increased log-gain variance scales with the exponent p. For example, increasing p from ½ to 1, doubles the log-gain. This statistical expectation-based gain approach is consistent with the contrast amplifications illustrated under the idealized FFDM acquisition model, while remaining applicable to situations where Beer’s Law does not strictly apply. As above, this suggests that increasing p should give larger ORs (or significant) in the logistic regression modeling.

### Spectral attributes of the chi-square images

The image c_m_(x, y) defined by Eq. [3] visually resembles a *mammogram* for sufficiently large m. However, its spectral properties differ from those of the mammogram used for its synthesis. Although these differences will not be investigated here in detail, we briefly show why this assertion holds. We take a Fourier transform approach, noting ensemble averaging in the image domain results in the same solution; this is to connect this derivation to related work referenced after the derivation. To illustrate the spectral differences, we use the Eq. [2] form without explicitly calculating its power spectrum. Using the convolution relationship, squaring the residual image, with m =1, and taking its two-dimensional Fourier transform gives:

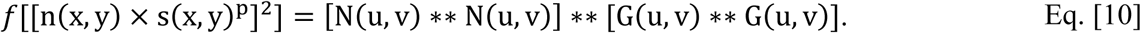

Here, *f*[·] is the 2D Fourier transform operator, g(x, y) = s(x,y)^p^, ∗∗ is the 2D convolution operator, capital letters indicate the Fourier transforms of the image factors, and (u, v) are two-dimensional Cartesian spatial frequency coordinates. Incorporating ensemble averaging with m >>1 yields key relationships: E[n(x, y)^2^] = 1 implies E[N(u, v) ∗∗ N(u, v)] = δ(u)δ(v). When G(u,v) is bandlimited to |u| or |v| ≤ F_c_, then H(u, v) = G(u, v) ∗∗ G(u, v) has broader spectral width than G(u, v) and is bandlimited to |u| or |v| ≤ 2F_c_. Substituting E[N(u, v) ∗∗ N(u, v)] = δ(u)δ(v) into Eq. [10] shows the ensemble averaged expectation yields H(u, v); the inverse Fourier transform of H(u, v) is s(x, y)^2p^. Consequently, in general G(u, v) ≠ H(u, v). The Fourier spectral properties of Eq. [10] will be investigated in the future. This derivation also used the artifact that the variation in the residual component in chi square image vanishes for large m. Notably, the expected ensemble averaged H(u, v) and resulting image domain s(x, y)^2p^ solutions outlined above are in agreement with a related Fourier domain formulation that used a single realization from Eq. [2] [43] (see supplementary material).

### Measurement standardization

We investigate various transformations applied to PBD measurements obtained under different settings to establish standardization. The objective is to evaluate the influence of a given transformation on logistic regression coefficients when merging measurements taken from different technologies, or populations, to increase the sample size. We assume if two or more populations are statistically similar, differences in their breast density distributions may arise from technological artifacts such as (1) image format differences, (2) digital image acquisition type, and (3) techniques used to produce the breast density measurements (applicable when using the same metric). Although a given method may produce significant results within multiple studies individually, merging measurements across studies that used different formatted image may induce unwanted variation in the model coefficients. We take a general approach to standardizing measurements. Importantly, this study is limited in what can be discerned from these modeling experiments directly. Conditional logistic regression is the appropriate methodology for evaluating the matched case-control design (described below): scale differences are accommodated, but distribution shifts may produce uncertainty, and site indicator coefficient estimates can be unstable.

Among the transformations investigated, an approximation to a probability integral transform (PIT) is the most general. A distribution is defined as the reference and other distributions are transformed to this reference. The reference distribution may be either external or internal: (1) a zero-mean unit-variance normal distribution is the external reference; and (2) the distribution from C-View image measurements is the internal reference (i.e., the most current technology used in this study). The PIT technique is more general than a linear transformation because it accommodates non-linear relationships, while reducing to a linear transformation when appropriate; no prior assumptions are imposed upon the functional form of the relationship. In any event, the approach should accommodate small sample sizes in the limiting situation. The PIT is investigated as it may be useful in future studies as well as for comparisons in this investigation. This technique is compared to the z-transformation, logarithmic transformation, and without a transformation, noting the logarithmic transformation is often used to reduce right-skewness and limit the influence of outliers. When applying the PIT to measures takne from small sample sizes (as in this present study), an optimized univariate kernel density estimation (KDE) technique is used to generate synthetic measurement samples. These additional samples are used to ensure integral approximations used for PIT computation are stable [49]. Incorporating KDE for small n yields an approximation for continuous PITs.

### Study Population

The new automated approach was applied to three successive case-control (CC) studies conducted over the course of a long-standing investigation, each defined by a given technology used to acquire its mammograms. Table 1 shows imaging technology for each study and image format details. Specifically, Study 1 encompassed 2D mammograms from a GE unit with 0.10mm pixel spacing; Study 2 encompassed 2D mammograms from Hologic units with 0.07mm pixel spacing; and Study 3 encompassed DBT datasets (volume and 2D synthetic images) from Hologic units with approximately 0.10mm pixel spacing. For brevity, we refer to these 3 studies as GE/Hologic raw-FFDM, GE/Hologic clinical-FFDM, or C-View (DBT), respectively. Although the Hologic and GE systems produce FFDM images, their detector technologies are different. Hologic units use direct x-ray conversion detector technology, whereas the GE unit used indirect x-ray conversion, producing different data representations. The parameter n was selected a priori based on the pixel spacing of a given image format to make the spatial averaging over approximately the same restricted local spatial scale: n = 4 (0.28mm) for Hologic FFDM; and n = 2 (0.20mm) for GE-FFDM and C-View images. For DBT images, pixel spacing varies by participant; we use 0.10 mm as an average approximation taken over all participants.

**Table 1.** Image format descriptions. Shown are each mammography unit’s manufacturer, pixel dynamic range, number of matched case-control pairs (n), and image type/usage (top row). Full field digital mammography (FFDM) formats involve case-control pairs with data available both for processing (raw) and for presentation. Hologic includes images acquired with Selenia FFDM and Dimensions digital breast tomosynthesis (DBT) units. The intended use for each type of format is provided parenthetically in the top row: “for processing” are raw mammograms not used for clinical purposes and “for presentation” are mammograms used for clinical purposes when considering FFDM.

| Image Format | FFDM<br>for processing<br>(raw) | FFDM<br>for presentation<br>(clinical) | DBT<br>C-View<br>(clinical) |
| --- | --- | --- | --- |
| General Electric | 14-bit<br>n = 180 | 12-bit<br>n = 180 | n/a<br>n/a |
| Hologic | 14-bit<br>n = 667 | 12-bit<br>n = 667 | 10-bit<br>n = 426 |

The participant selection protocol was the same for each of the three studies, described previously [50, 51]. All procedures were carried out in accordance with the relevant guidelines and regulations approved by the Institutional Review Board of the University of South Florida, Tampa, FL under protocol 104715_CR000002. These studies were designed specifically to investigate image measures that could be used for risk prediction modeling. To isolate the effects of image measurements, we selected controls that were matched individually to cases on key variables known to strongly influence breast physiology i.e., age within ± 2 years, hormone replacement (HR) usage (ever or never) and HR duration intervals (± 2 years). Controls were matched to cases on mammography units and screening history as well. All cases had pathologically confirmed unilateral breast cancer. The unaffected breast was analyzed with PD_a_ for cases and the corresponding latterly matched breast of the control was analyzed, forming the respective matched case-control pair. Only craniocaudal views were considered to avoid chest wall interference.

When this institution’s mammography centers were transitioning to Hologic-DBT from Hologic-FFDM, the HD Combo Mode acquisition was used as standard of care imaging when operating DBT units. In this mode, two-dimensional and DBT acquisitions are taken in sequence without repositioning the patient or changing the breast compression. A total of 78 CC pairs underwent mammography during this transition period straddling studies 2 and 3 and had both DBT acquisitions and 2D FFDM images for analysis. For the present investigation, to optimize numbers of independent observations in each of the defined studies and weighting toward the newer DBT technology, 78 CC pairs were included in the present analysis with Study 3; hence, data were analyzed for a total of 180, 241 and 426 CC pairs in studies 1, 2 and 3, respectively.

### Risk prediction modeling

To identify the optimal operating parameters for PD_a_ within the scope of this investigation, OR tables were constructed for each image format as functions of (1) ensemble averaging m, (2) detection search window size n (in pixel units), (3) two chi-square detection parameters governing the search window operations, and (4) the exponent p. Odds ratios and corresponding regression coefficients were determined with conditional logistic regression. Odds ratios with CIs and the corresponding intrinsic PBD intrinsic distribution’s mean and standard deviation are cited in each cell of these tables. These tables are relatively large (64 cells), and model parameters were generated after taking the natural logarithm of the PBD measurements, thereby giving another transformation comparison. In all models, we controlled for body mass index (BMI) and ethnicity; age and hormonal attributes were matching factors in case-control selections.

For large, automated batch processing tasks, such as those used in this study, working in the logarithmic domain is our standard approach when investigating image measures because their intrinsic distributions may be positive-skewed (i.e., normally no chance of a negative valued finding). This step often gives more accurate estimates but can compress the intrinsic distribution to a point where a unit shift may span most of its width (spanning left to right tails), artificially inflating both the ORs and regression coefficients. Although the intrinsic distribution summaries are cited in each OR cell, the reported modeling results are based on the logarithmic transformed PBD measurements; ORs were generated in per standard deviation shift of the transformed distribution.

To assist in making algorithm parameter choices from the OR tables, historical PBD measurements from the Cumulus technique were used as reference. These are tabulated in Table 2 and were distilled from these studies: (1 and 2) are from [3]; (3) is from [47]; (4) is from [4]; (5 and 6) are from [52]; and (7 and 8) are from [53]. This list includes measurements from GE-FFDM, Hologic-FFDM, and digitized film mammograms from three different types of digitizers. For modeling illustration, a single PD_a_ detection output distribution was selected from each image format; when significant, selection is based on similarity with the historical PBD measurements via mean and standard deviation comparisons. Averaging the means and standard deviations separately reported in Table 2 yields the approximate PBD historical references: µ ≈ 25 and σ ≈ 17 used here. Operating parameters near these reference values are used for the primary illustrations.

**Table 2.** Historical percentage of breast density measurement distribution summaries: a brief survey of prior percentage of breast density (PBD) distribution summaries are given mean (µ) and standard deviation (σ). The unit or detector specifics and participant numbers (n) are also provided. PBD measurements were made with the user-assisted Cumulus method. Numbers 4-6 reference film digitizers. Specific study citations are identified in a subsection within the Methods Section titled *Risk prediction modeling*.

| Unit / detector type | n | Image type | $\mu$ | $\sigma$ |
| --- | --- | --- | --- | --- |
| 1.Senographe 2000D | 360 | raw-FFDM | 19.5 | 15.2 |
| 2.Senographe 2000D | 360 | clinical-FFDM | 21.1 | 13.7 |
| 3.Selenia & Dimensions | 638 | clinical-FFDM | 25.3 | 11.6 |
| 4.Kodak Lumiscan 75 | 712 | digitized film | 26.5 | 14.6 |
| 5.Lumysis 85 and<br>VIDAR CAD PRO<br>Advantage | 1900 | digitized film | 37.6 | 19.8 |
| 6.Lumysis 85 and<br>VIDAR CAD PRO<br>Advantage | 3921 | digitized film | 32.1 | 19.6 |
| 7.Hologic Selenia | 550 | raw-FFDM | 14.1 | 12.4 |
| 8.Hologic Selenia | 550 | clinical-FFDM | 17.4 | 10.1 |

### Frequently used abbreviations and conventions

The following abbreviations and conventions apply throughout this report: artificial intelligence (AI); case-control (CC); digital breast tomosynthesis (DBT); full field digital mammography (FFDM); General Electric (GE); odds ratio (OR); signal-dependent noise (SDN). A normalized histogram is referred to as an empirical probability density function (pdf). When a pdf is made from a given measurement without a transformation, it is referred to as the intrinsic pdf. Intrinsic is used in the same sense for other quantities as well. The interger parameter n is used to define both the number of CC pairs and the detention search window size (context is clear). A standardized normal distribution has zero-mean with unit variance. C-View images are two-dimensional images, constructed from DBT acquisitions, and are used for clinical purposes. DBT provides a limited angle quasi three-dimensional images reconstructed breast volume. In the DICOM header fields, images labeled *for presentation* are used for clinical purposes and those labeled *for processing* are loosely raw images. Clinical two-dimensional FFDM images are created from the respective raw images with vendor proprietary image processing algorithms. Many facilities destroy the raw images. The various images have different data representations, which are referred to as format differences.

## Results

### Image format differences and breast density detection examples

Figure 1, top-row shows mammograms acquired at the same time point for two women in different formats: (1) panes 1-3 show three images from woman-1, and (2) panes 4-5 show two images from woman-2. These are used to exemplify the approach and illustrate natural variations throughout this work. Pane-1 shows a Hologic raw-FFDM image after inverting its intensities, pane-2 shows the corresponding clinical FFDM-image, pane-3 shows a C-View image from Hologic DBT (panes 1-3 show images from woman-1 at the same time point), pane-4 shows a GE raw-FFDM image after inverting its intensities, and pane-5 shows the associated GE clinical FFDM-image (both GE images are from woman-2 at the same time point). These women were selected at random from the respective studies. These multiple images from a given woman are all valid image formats. The breast density intra-variation within each woman’s images is clear visually. The second row shows the respective chi-square images derived with varying degrees of ensemble averaging: from left to right m = (25, 5, 50, 25, 100). As mentioned previously, images in the middle row of this figure visually resemble mammograms. The corresponding PD_a_ images are shown in the bottom row with these results: (1) 13.2%; (2) 23.8%; (3) 25%; (4) 36.3%; and (5) 52.0% respectively. Each of these percentages were derived from a specific set of detection parameters discussed below. These demonstrate the scale variation problem; within each format, the CC distributions for PD_a_ produced significant ORs, although PBD clearly varies across formats for the same woman due to the differences. We are left with the obvious quandary of choosing the best representation, assuming one exists.

### Detection parameters and modeling results

The detection parameters selected for PD_a_ are discussed in the context of the modeling results. Historical distribution summaries were used in conjunction with significant OR findings. Because the CC studies used here were designed for image measurement investigations, one guiding premise was that significant ORs provide an objective truth given breast density is a significant risk factor. The degree of ensemble averaging (m) for each format was determined through initial experimentation (not shown here). The spatial averaging extent was set a priori by considering the intrinsic pixel spacing with n = 4 for the higher resolution images (i.e., Hologic FFDM images) and n = 2 for the other image formats; this ensures we analyzed consistent fine spatial scale variations over all formats. Initial experimentation indicated that setting the exponent p = 1 provided superior results (more cells with significant ORs). Accordingly, OR tables for p = ½ are not shown. The respective OR findings are shown in tables 3-7. Odds ratios are provided in each cell with confidence intervals as a function of the two detection parameters. Although ORs were generated by first log-transforming the intrinsic distributions, reported means and standard deviations (i.e., summaries) in these cells are derived from the intrinsic distributions. Blue shaded cells in each table illustrate operating parameters that provided significant ORs that have distribution summaries approximately within the range of the historical references. These were used as an approximate objective truth because other cells with relative outlier summaries with significant ORs may be valid as well; these other cells also provide examples of the scaling variations in the PD_a_ distributions both within and across formats. The objective of the optimization procedure was to adjust the four parameters to optimize the number of possible acceptable operating points for a given format. Importantly, these results require special attention: (1) all cells for Hologic-raw-FFDM, C-View and, GE clinical-FFDM have significant ORs; and (2) most cells for the Hologic clinical-FFDM have significant ORs. In each table, the cell denoted with bold fonts and an asterisk defines the respective operating points selected for this investigation; these parameters are stable choices because perturbing them slightly produced similar findings (surrounding cells for the most part). In summary, significant ORs differed by image format (standard deviation shift): (1) ORs ∼ 1.31-1.60 derived from images used for clinical purposes; and ORs ∼ 1.20 – 1.31 derived from raw FFDM images. The tables for p = ½ are not shown.

**Table 3.** General Electric raw-FFDM odds ratio table: this gives the odd ratios (ORs) for PD_a_ with m =25 (number of ensemble averages) and n= 2 (detection search window size in pixels) over the range of detection parameters; the top row gives detection parameter 1 and the first column detection parameter 2. There are three metrics cited in each cell: (1) distribution mean (µ, top); (2) distribution standard deviation (σ, middle); and (3) an OR with confidence intervals parenthetically. Blue shading indicates possible operating parameters for PDa. The cell with bold fonts and asterisk gives the detection parameters used throughout the report’s examples.

| Detection Parameters | $1^{-1}$ | $1^{-2}$ | $1^{-3}$ | $1^{-4}$ | $1^{-5}$ | $1^{-6}$ | $1^{-7}$ | $1^{-8}$ | |
| --- | --- | --- | --- | --- | --- | --- | --- | --- | --- |
| $1^{-1}$ | 49.5<br>6.3<br>1.04 (0.83, 1.30) | 41.8<br>10.3<br>1.12 (0.89, 1.40) | 36.1<br>12.3<br>1.15 (0.92, 1.44) | 31.7<br>13.3<br>1.18 (0.94, 1.48) | 28.2<br>13.6<br>1.20 (0.96, 1.51) | 25.2<br>13.7<br>1.22 (0.97, 1.53) | 22.7<br>13.6<br>1.23 (0.98, 1.55) | 3.6<br>6.3<br>n/a | $\mu$<br>$\sigma$<br>OR |
| $1^{-2}$ | 43.5<br>7.3<br>1.12 (0.90, 1.40) | 34.8<br>10.8<br>1.19 (0.95, 1.49) | 28.7<br>12.2<br>1.22 (0.97, 1.54) | 24.6<br>12.6<br>1.25 (0.99, 1.58) | 21.3<br>12.7<br>1.27 (1.01, 1.61) | 18.7<br>12.5<br>1.29 (1.02, 1.63) | 16.6<br>12.2<br>1.20 (1.03, 1.66) | 2.2<br>4.8<br>n/a | $\mu$<br>$\sigma$<br>OR |
| $1^{-3}$ | 39.8<br>7.4<br>1.16 (0.93, 1.45) | 30.7<br>10.5<br>1.22 (0.97, 1.54) | 24.8<br>11.6<br>1.26 (1.00, 1.59) | **20.8\***<br>**11.9**<br>**1.29 (1.02, 1.63)** | 17.8<br>11.7<br>1.31 (1.03, 1.66) | 15.4<br>11.4<br>1.32 (1.04, 1.68) | 13.5<br>11.0<br>1.34 (1.05, 1.70) | 1.6<br>3.9<br>n/a | $\mu$<br>$\sigma$<br>OR |
| $1^{-4}$ | 37.2<br>7.1<br>1.17 (0.94, 1.46) | 28.0<br>10.1<br>1.24 (0.98, 1.56) | 22.3<br>11.0<br>1.28 (1.01, 1.62) | 18.4<br>11.1<br>1.30 (1.03, 1.65) | 15.5<br>10.9<br>1.32 (1.04, 1.68) | 13.4<br>10.5<br>1.34 (1.05, 1.71) | 11.6<br>10.1<br>1.35 (1.06, 1.72) | 1.2<br>3.2<br>n/a | $\mu$<br>$\sigma$<br>OR |
| $1^{-5}$ | 35.4<br>6.7<br>1.17 (0.94, 1.46) | 26.1<br>9.5<br>1.24 (0.99, 1.57) | 20.46<br>10.4<br>1.29 (1.02, 1.63) | 16.7<br>10.4<br>1.31 (1.04, 1.67) | 14.0<br>10.2<br>1.34 (1.05, 1.70) | 12.0<br>9.8<br>1.35 (1.06, 1.72) | 10.3<br>9.4<br>1.37 (1.07, 1.74) | 0.9<br>2.6<br>n/a | $\mu$<br>$\sigma$<br>OR |
| $1^{-6}$ | 34.1<br>6.4<br>1.17 (0.93, 1.46) | 24.7<br>9.0<br>1.25 (0.99, 1.57) | 19.2<br>9.8<br>1.29 (1.02, 1.64) | 15.5<br>9.8<br>1.32 (1.04, 1.68) | 12.9<br>9.6<br>1.34 (1.06, 1.71) | 10.9<br>9.2<br>1.36 (1.07, 1.73) | 9.<br>48.7<br>1.37 (1.08, 1.75) | 0.73<br>2.2<br>n/a | $\mu$<br>$\sigma$<br>OR |
| $1^{-7}$ | 33.0<br>6.09<br>1.16 (0.93, 1.45) | 23.6<br>8.6<br>1.25 (0.99, 1.57) | 18.1<br>9.3<br>1.29 (1.02, 1.64) | 14.5<br>9.3<br>1.33 (1.04, 1.69) | 12.0<br>9.00<br>1.35 (1.06, 1.72) | 10.q<br>8.6<br>1.37 (1.07, 1.74) | 8.6<br>8.1<br>1.38 (1.08, 1.76) | 0.59<br>1.9<br>n/a | $\mu$<br>$\sigma$<br>OR |
| $1^{-8}$ | 28.4<br>4.9<br>1.01 (0.81, 1.26) | 19.2<br>5.9<br>1.20 (0.95, 1.50) | 14.0<br>6.2<br>1.28 (1.01, 1.62) | 10.5<br>6.0<br>1.33 (1.04, 1.69) | 8.2<br>5.4<br>1.36 (1.07, 1.73) | 6.5<br>4.9<br>1.38 (1.08, 1.76) | 5.2<br>4.4<br>1.40 (1.10, 1.79) | 0.08<br>0.53<br>n/a | $\mu$<br>$\sigma$<br>OR |

**Table 4.** General Electric clinical-FFDM odds ratio table: this gives the odd ratios (ORs) for PD_a_ with m =100 (number of ensemble averages) and n = 2 (detection search window size in pixels) over the range of detection parameters; the top row gives detection parameter 1 and the first column detection parameter 2. There are three metrics cited in each cell: (1) distribution mean (µ, top); (2) distribution standard deviation (σ, middle); and (3) an OR with confidence intervals parenthetically. Blue shading indicates possible operating parameters for PDa. The cell with bold fonts asterisk give the detection parameters used throughout this report’s examples.

| Detection Parameters | $1^{-1}$ | $1^{-2}$ | $1^{-3}$ | $1^{-4}$ | $1^{-5}$ | $1^{-6}$ | $1^{-7}$ | $1^{-8}$ | |
| --- | --- | --- | --- | --- | --- | --- | --- | --- | --- |
| $1^{-1}$ | 51.7<br>9.7<br>1.62 (1.26, 2.08) | 33.9<br>11.9<br>1.63 (1.26, 2.11) | **24.0\***<br>**12.1**<br>**1.62 (1.25, 2.10)** | 17.9<br>11.5<br>1.61 (1.24, 2.09) | 13.9<br>10.6<br>1.61 (1.24, 2.09) | 11.0<br>9.7<br>1.61 (1.24, 2.08) | 8.8<br>8.8<br>1.60 (1.24, 2.07) | 0.5<br>1.3<br>n/a | $\mu$<br>$\sigma$<br>OR |
| $1^{-2}$ | 42.3<br>9.4<br>1.65 (1.28, 2.13) | 26.4<br>11.2<br>1.62 (1.25, 2.10) | 18.3<br>11.0<br>1.60 (1.24, 2.08) | 13.5<br>10.1<br>1.60 (1.23, 2.07) | 10.4<br>9.1<br>1.59 (1.22, 2.06) | 8.1<br>8.2<br>1.58 (1.22, 2.04) | 6.5<br>7.3<br>1.56 (1.21, 2.02) | 0.3<br>0.9<br>n/a | $\mu$<br>$\sigma$<br>OR |
| $1^{-3}$ | 37.9<br>8.8<br>1.65 (1.28, 2.14) | 23.0<br>10.4<br>1.61 (1.24, 2.09) | 15.8<br>10.0<br>1.59 (1.23, 2.07) | 11.5<br>9.1<br>1.58 (1.22, 2.05) | 8.7<br>8.1<br>1.57 (1.21, 2.04) | 6.8<br>7.2<br>1.56 (1.20, 2.02) | 5.3<br>6.3<br>1.54 (1.19, 1.99) | 0.2<br>0.6<br>n/a | $\mu$<br>$\sigma$<br>OR |
| $1^{-4}$ | 35.3<br>8.1<br>1.65 (1.28, 2.14) | 21.1<br>9.6<br>1.61 (1.24, 2.08) | 14.3<br>9.2<br>1.58 (1.22, 2.05) | 10.3<br>8.2<br>1.57 (1.21, 2.04) | 7.8<br>7.3<br>1.56 (1.21, 2.02) | 6.0<br>6.4<br>1.54 (1.19, 1.99) | 4.6<br>5.6<br>1.52 (1.18, 1.97) | 0.2<br>0.5<br>n/a | $\mu$<br>$\sigma$<br>OR |
| $1^{-5}$ | 33.6<br>7.4<br>1.65 (1.28, 2.14) | 19.8<br>8.9<br>1.60 (1.24, 2.07) | 13.3<br>8.5<br>1.58 (1.22, 2.05) | 9.5<br>7.6<br>1.57 (1.21, 2.03) | 7.1<br>6.7<br>1.55 (1.20, 2.01) | 5.4<br>5.8<br>1.54 (1.19, 1.98) | 4.1<br>5.0<br>1.51 (1.17, 1.95) | 0.2<br>0.4<br>n/a | $\mu$<br>$\sigma$<br>OR |
| $1^{-6}$ | 32.5<br>6.9<br>1.65 (1.28, 2.14) | 19.0<br>8.4<br>1.60 (1.23, 2.07) | 12.6<br>7.9<br>1.58 (1.22, 2.04) | 8.9<br>7.1<br>1.56 (1.21, 2.02) | 6.6<br>6.1<br>1.55 (1.20, 2.00) | 5.0<br>5.3<br>1.53 (1.19, 1.98) | 3.8<br>4.5<br>1.51 (1.17, 1.95) | 0.1<br>0.3<br>n/a | $\mu$<br>$\sigma$<br>OR |
| $1^{-7}$ | 31.6<br>6.5<br>1.65 (1.28, 2.14) | 18.3<br>7.9<br>1.60 (1.23, 2.07) | 12.1<br>7.5<br>1.58 (1.22, 2.04) | 8.5<br>6.6<br>1.56 (1.21, 2.02) | 6.2<br>5.7<br>1.55 (1.20, 2.00) | 4.7<br>4.9<br>1.53 (1.19, 1.97) | 3.6<br>4.1<br>1.51 (1.17, 1.94) | 0.1<br>0.3<br>n/a | $\mu$<br>$\sigma$<br>OR |
| $1^{-8}$ | 29.1<br>5.2<br>1.62 (1.26, 2.10) | 16.4<br>6.3<br>1.60 (1.23, 2.08) | 10.5<br>5.8<br>1.58 (1.22, 2.05) | 7.2<br>5.0<br>1.57 (1.21, 2.03) | 5.1<br>4.1<br>1.56 (1.20, 2.01) | 3.7<br>3.3<br>1.54 (1.19, 1.98) | 2.7<br>2.7<br>1.51 (1.17, 1.95) | 0.1<br>0.2<br>n/a | $\mu$<br>$\sigma$<br>OR |

**Table 5.** Hologic raw-FFDM odds ratio table: this gives the odd ratios (ORs) for PD_a_ with m = 25 (number of ensemble averages) and n= 4 (detection search window size in pixels) over the range of detection parameters; the top row gives detection parameter 1 and the first column detection parameter 2. There are three metrics cited in each cell: (1) distribution mean (µ, top); (2) distribution standard deviation (σ, middle); and (3) an OR with confidence intervals parenthetically. Blue shading indicates possible operating parameters for PDa. The cell with bold fonts and asterisk gives the detection parameters used throughout this report’s examples.

| Detection Parameters | $1^{-1}$ | $1^{-2}$ | $1^{-3}$ | $1^{-4}$ | $1^{-5}$ | $1^{-6}$ | $1^{-7}$ | $1^{-8}$ | |
| --- | --- | --- | --- | --- | --- | --- | --- | --- | --- |
| $1^{-1}$ | 49.3<br>10.79<br>1.21 (1.07, 1.37) | 38.9<br>15.8<br>1.23 (1.08, 1.39) | 32.0<br>17.7<br>1.24 (1.09, 1.41) | 27.0<br>18.4<br>1.25 (1.10, 1.42) | 23.3<br>18.4<br>1.25 (1.10, 1.43) | 20.4<br>18.1<br>1.26 (1.11, 1.43) | 18.0<br>17.6<br>1.26 (1.11, 1.44) | 4.6<br>9.7<br>n/a | $\mu$<br>$\sigma$<br>OR |
| $1^{-2}$ | 42.4<br>12.0<br>1.25 (1.10, 1.42) | 31.6<br>16.0<br>1.26 (1.11, 1.43) | **25.0\***<br>**17.0**<br>**1.26 (1.11, 1.44)** | 20.6<br>16.9<br>1.27 (1.12, 1.44) | 17.5<br>16.5<br>1.27 (1.12, 1.45) | 15.1<br>15.9<br>1.28 (1.12, 1.45) | 13.2<br>15.3<br>1.28 (1.12, 1.45) | 3.1<br>7.6<br>n/a | $\mu$<br>$\sigma$<br>OR |
| $1^{-3}$ | 38.6<br>11.9<br>1.27 (1.11, 1.44) | 27.7<br>15.3<br>1.27 (1.12, 1.45) | 21.4<br>15.8<br>1.28 (1.12, 1.46) | 17.3<br>15.5<br>1.28 (1.13, 1.46) | 14.5<br>14.9<br>1.29 (1.13, 1.46) | 12.4<br>14.3<br>1.29 (1.13, 1.46) | 10.7<br>13.5<br>1.28 (1.13, 1.46) | 2.4<br>6.3<br>n/a | $\mu$<br>$\sigma$<br>OR |
| $1^{-4}$ | 36.2<br>11.4<br>1.28 (1.12, 1.45) | 25.3<br>14.4<br>1.28 (1.13, 1.46) | 19.1<br>14.8<br>1.29 (1.13, 1.47) | 15.3<br>14.4<br>1.29 (1.13, 1.47) | 12.6<br>13.7<br>1.29 (1.13, 1.47) | 10.7<br>13.0<br>1.29 (1.13, 1.47) | 9.2<br>12.3<br>1.28 (1.13, 1.46) | 2.0<br>5.6<br>n/a | $\mu$<br>$\sigma$<br>OR |
| $1^{-5}$ | 34.7<br>10.9<br>1.28 (1.13, 1.46) | 23.7<br>13.7<br>1.29 (1.13, 1.47) | 17.6<br>13.9<br>1.29 (1.14, 1.47) | 13.9<br>13.4<br>1.29 (1.14, 1.47) | 11.4<br>12.7<br>1.30 (1.14, 1.47) | 9.5<br>12.0<br>1.29 (1.14, 1.47) | 8.1<br>11.3<br>1.28 (1.13, 1.46) | 1.7<br>5.0<br>n/a | $\mu$<br>$\sigma$<br>OR |
| $1^{-6}$ | 33.6<br>10.4<br>1.29 (1.13, 1.46) | 22.5<br>13.0<br>1.29 (1.14, 1.47) | 16.6<br>13.2<br>1.30 (1.14, 1.48) | 12.9<br>12.6<br>1.30 (1.14, 1.48) | 10.5<br>11.9<br>1.30 (1.14, 1.48) | 8.7<br>11.2<br>1.29 (1.14, 1.47) | 7.4<br>10.5<br>1.28 (1.13, 1.46) | 1.5<br>4.6<br>n/a | $\mu$<br>$\sigma$<br>OR |
| $1^{-7}$ | 32.7<br>9.9<br>1.29 (1.13, 1.46) | 21.7<br>12.4<br>1.30 (1.14, 1.47) | 15.7<br>12.5<br>1.30 (1.14, 1.48) | 12.1<br>11.9<br>1.30 (1.14, 1.48) | 9.8<br>11.2<br>1.30 (1.14, 1.48) | 8.1<br>10.6<br>1.29 (1.14, 1.47) | 6.8<br>9.9<br>1.28 (1.13, 1.46) | 1.3<br>4.2<br>n/a | $\mu$<br>$\sigma$<br>OR |
| $1^{-8}$ | 29.5<br>7.7<br>1.24 (1.09, 1.40) | 18.5<br>9.3<br>1.28 (1.12, 1.46) | 12.7<br>9.0<br>1.29 (1.13, 1.47) | 9.3<br>8.2<br>1.29 (1.14, 1.47) | 7.1<br>7.5<br>1.29 (1.14, 1.47) | 5.5<br>6.7<br>1.29 (1.13, 1.47) | 4.4<br>6.1<br>1.28 (1.12, 1.45) | 0.5<br>2.0<br>n/a | $\mu$<br>$\sigma$<br>OR |

### Signal-dependent noise characteristic evaluation

In this section, we discuss why synthesizing the SDN to a specified quadratic structural form constitutes the major innovation responsible for producing significant results from images in different formats. This formulation may be defined within a more general theoretical framework. For each woman, the ideal s(x, y) was approximated, and multiple realizations of the squared residual image were then synthesized, treating the model as a two-dimensional stochastic process conditioned on this approximation. For p = 1, we expect the noise variance to exhibit a quadratic statistical relationship with the mean signal when analyzing the squared and ensemble averaged residual image c_m_(x, y) in Eq. [3]. The intrinsic SDN structure for the three types of clinical mammograms shown in the top row of Figure 1 (panes 2, 3, and 5 corresponding with Hologic-clinical, Hologic C-view, and GE-clinical respectively) are displayed in the top row of Figure 2, respectively. These show the noise variance (y-axis) as a function of the local pixel average (x-axis) in each mammogram. Potential problems for the automated detection can be noted in each example. Both the Hologic-clinical and C-View plots are concave-down with first derivatives that have maximums over the signal-average range. Here, two average signal values correspond with the same noise variance. When detection thresholds are located between horizontal line intersects, it will cause algorithm confusion. The GE-clinical plot has approximately two increasing *linear* sections connected with an elbow, which is also less than optimal. The corresponding empirical plots derived from c_m_(x, y) in Eq. [3] are shown in the bottom row of Figure 2. In each pane, the plot computed from the data is shown in black (empirically derived curves have been smoothed for illustration purposes, suppressing natural variation) and is compared with the fitted curve using a quadratic polynomial model shown in red. These modeled curves show close agreement with the empirical curves, as predicted. Similar agreement was noted across all images and formats (not shown). Such predictable relationships derived from different formats, both within and across, stabilized the density detection algorithm; the quadratic structure provided sufficient contrast between the variances of the two tissue types and produced many cells with significant ORs within and across image formats. Moreover, these findings agree with the variance-gain derivations, linking either imaging physics or imaging processing frameworks to the significance of the exponent choice. The same results provide an explanation for why the algorithm-performance was not optimal in both past experiments and in this work when the local SDN structure obeyed Poisson statistics (i.e., when p = ½ was applicable) as applicable to raw FFDM images.

**Figure 1.**
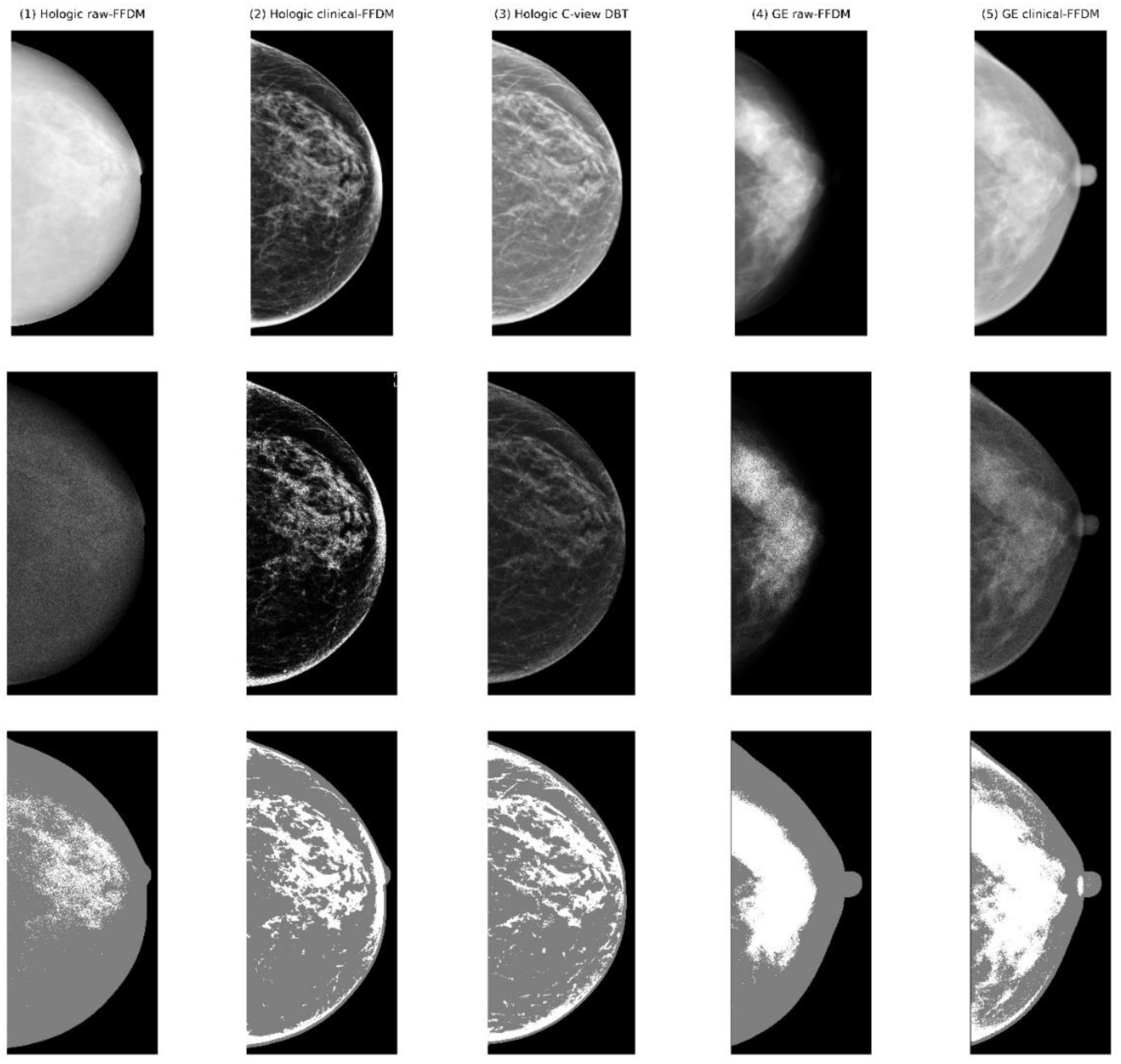
Image format differences and corresponding PD_a_ examples: in the top row (left to right) panes 1-3 show mammograms from woman-1, acquired *simultaneously* in three different formats, and panes 3-4 show mammograms from women-2 acquired *simultaneously* in two different formats: (1) woman-1, Hologic raw FFDM; (2) woman-1, Hologic clinical FFDM; (3) woman-1, Hologic C-View, (4-5) woman-2, GE raw FFDM and GE clinical FFDM images, respectively. The middle row shows the corresponding cm(x, y) images from Eq. [3] obtained with m ensemble averages followed n×n spatial averaging: (1) m = 25 and n = 4; (2) m = 5 and n = 4; (3) m = 50 and n = 2; (4) m = 25 and n = 2; and (5) m = 100 and n = 2. The bottom row shows the adipose and dense areas detected by PDa: (1) 13.2%; (2) 23.8%; (3) 25%; (4) 36.3%; and (5) 52.0%. Images shown in the middle row were adjusted for viewing purposes and all have the same window level and width.

**Figure 2.**
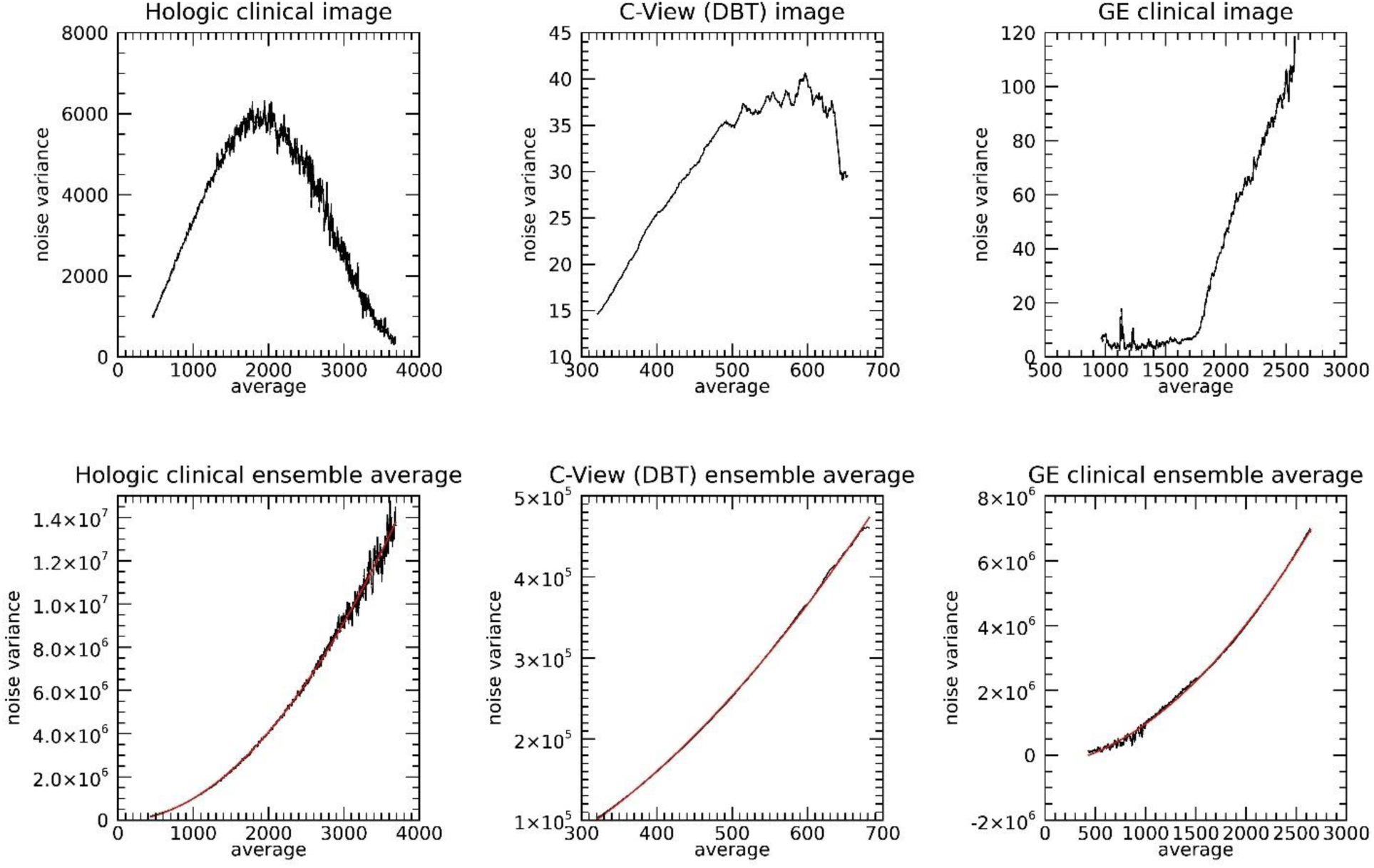
Signal-dependent noise variance plot examples: these plots show expected pixel values (average) on the x-axes and their expected variances on the y-axes. The top row shows intrinsic plots derived from images used for clinical purposes shown in Figure 1: Hologic-clinical FFDM; Hologic C-View from DBT; and GE-clinical FFDM. The bottom row shows the respective plots derived from the chi-square images, also shown in Figure 1: black curves show data and red shows the fitted quadratic-model curves derived from curve fitting analysis.

A comment is warranted to qualify the selection of the exponent p and the ensemble parameter m. We cannot claim p has been globally optimized because only two values were considered. However, the quadratic choice produced may benefits explained above. Likewise, we cannot claim the choice for m was optimal for each format; it was selected by examining the same sample range of specific values for each image format within the chi-square detection parameters (examining number of cells with significant ORs) and visual observation, where detection results fail to resemble the respective image density. Also, increasing m beyond a practical limit will attenuate the local variation induced by the noise modulation, which will negatively impact the detection performance as discussed below.

### Breast density measurement standardization

The main standardization and transformation results from PD_a_ are discussed first followed by alternative experiments. Table 8 shows the results for mammograms used clinically. The number of CC pairs used in the respective analysis are reported in the uppermost row with the headings. The total n reported that the last column is derived by combining unique pairs from each image format: n = 180 + 241 + 426 = 847 unique pairs. Below the heading row, this table is organized with three rows (i.e., top middle and bottom) each with multiple findings that are numbered. The regression parameter β for each type of transformation is cited in the top-row. Here (1 and 2) are best compared, and (3 and 4) are best compared. The largest impact is found within the GE clinical-FFDM findings. We posit the GE findings are due to small sample size uncertainty and measurement distribution differences (discussed below). There were small changes in these coefficients when considering the PIT (comparing 1 with 2) or z-score transformation (comparing 3 with 4). In the middle row, the respective ORs are reported in unit shift [again, (1 and 2) are best compared, and (3 with 4) are best compared]. Within these comparison groups, the respective ORs in unit shift were similar within and across formats. In the bottom row, ORs are reported in standard deviation shift and were similar within formats (comparing 1-5); excluding the GE format, ORs were similar within and across formats confined to the comparison groups. Similarities in the last (total) column findings may be driven by the relatively large n in the C-View format, which is investigated further below with a specific example. Across formats, ORs in unit shift varied from 1.05 - 1.07, exhibiting relative consistency when considering the image format differences. The bottom row shows the ORs in per standard deviation shift. Within format, these varied slightly over the transformation type. Across formats, the OR from the GE format was the largest, again probably due to sampling uncertainty. For the other formats, the findings were similar when considering the CIs. Sampling uncertainly is illustrated by comparing the respective ORs between total Hologic column (n = 667 pairs) with the corresponding smaller sample column (n = 241 pairs). The combined column indicates PD_a_ can be combined to form a larger case-control study with n = 847 pairs. The similarities between the PIT approximations and z-score findings are an indication that the respective PITs have a strong linear trend.

**Table 6.** Hologic clinical-FFDM odds ratio table: this gives the odd ratios (ORs) for PD_a_ with m = 5 (number of ensemble averages) and n= 4 (detection search window size in pixels) over the range of detection parameters; the top row gives detection parameter 1 and the first column detection parameter 2. There are three metrics cited in each cell: (1) distribution mean (µ, top); (2) distribution standard deviation (σ, middle); and (3) an OR with confidence intervals parenthetically. Blue shading indicates possible operating parameters for PDa. The cell with bold fonts and asterisk gives the detection parameters used throughout this report’s examples.

| Detection Parameters | $1^{-1}$ | $1^{-2}$ | $1^{-3}$ | $1^{-4}$ | $1^{-5}$ | $1^{-6}$ | $1^{-7}$ | $1^{-8}$ | |
| --- | --- | --- | --- | --- | --- | --- | --- | --- | --- |
| $1^{-1}$ | 43.0 | 37.3 | 33.4 | 30.5 | 28.1 | 26.1 | 24.4 | 19.9 | $\mu$ |
| | 5.1 | 6.1 | 6.7 | 7.2 | 7.5 | 7.7 | 7.9 | 8.0 | $\sigma$ |
|  | 1.03 (0.90, 1.17) | 1.12 (0.98, 1.28) | 1.17 (1.02, 1.35) | 1.21 (1.05, 1.39) | 1.23 (1.07, 1.42) | 1.25 (1.08, 1.44) | 1.26 (1.10, 1.46) | 1.29 (1.12, 1.49) | OR |
| $1^{-2}$ | 40.4 | 34.7 | 30.9 | 28.1 | 25.8 | 23.9 | 22.3 | 18.0 | $\mu$ |
| | 4.9 | 5.9 | 6.6 | 7.0 | 7.3 | 7.5 | 7.6 | 7.6 | $\sigma$ |
|  | 1.07 (0.94, 1.23) | 1.17 (1.02, 1.34) | 1.22 (1.06, 1.40) | 1.25 (1.08, 1.43) | 1.26 (1.10, 1.46) | 1.28 (1.11, 1.47) | 1.29 (1.12, 1.48) | 1.31 (1.14, 1.50) | OR |
| $1^{-3}$ | 38.6 | 33.0 | 29.3 | 26.5 | 24.3 | 22.5 | 20.9 | 16.8 | $\mu$ |
| | 4.7 | 5.8 | 6.4 | 6.8 | 7.1 | 7.3 | 7.4 | 7.3 | $\sigma$ |
|  | 1.11 (0.97, 1.27) | 1.20 (1.04, 1.38) | 1.24 (1.08, 1.43) | 1.27 (1.10, 1.46) | 1.28 (1.11, 1.48) | 1.29 (1.12, 1.49) | 1.30 (1.13, 1.49) | 1.31 (1.14, 1.51) | OR |
| $1^{-4}$ | 37.3 | 31.7 | 28.0 | 25.3 | 23.2 | 21.4 | 19.8 | 15.9 | $\mu$ |
| | 4.6 | 5.6 | 6.3 | 6.7 | 6.9 | 7.1 | 7.1 | 7.0 | $\sigma$ |
|  | 1.13 (0.98, 1.29) | 1.22 (1.06, 1.40) | 1.26 (1.09, 1.45) | 1.28 (1.11, 1.47) | 1.29 (1.12, 1.49) | 1.30 (1.13, 1.49) | 1.30 (1.13, 1.50) | 1.31 (1.14, 1.51) | OR |
| $1^{-5}$ | 36.2 | 30.6 | 27.0 | **24.4\*** | 22.2 | 20.5 | 19.0 | 15.1 | $\mu$ |
| | 4.4 | 5.5 | 6.1 | **6.5** | 6.7 | 6.9 | 6.9 | 6.8 | $\sigma$ |
|  | 1.14 (0.99, 1.31) | 1.23 (1.07, 1.42) | 1.27 (1.10, 1.46) | **1.29 (1.12, 1.48)** | 1.30 (1.13, 1.49) | 1.30 (1.13, 1.50) | 1.31 (1.14, 1.50) | 1.32 (1.15, 1.51) | OR |
| $1^{-6}$ | 35.3 | 29.8 | 26.2 | 23.6 | 21.5 | 19.8 | 18.3 | 14.5 | $\mu$ |
| | 4.3 | 5.4 | 6.0 | 6.3 | 6.6 | 6.7 | 6.7 | 6.6 | $\sigma$ |
|  | 1.15 (1.00, 1.32) | 1.24 (1.08, 1.43) | 1.27 (1.10, 1.47) | 1.29 (1.12, 1.48) | 1.30 (1.13, 1.50) | 1.31 (1.14, 1.50) | 1.31 (1.14, 1.51) | 1.32 (1.15, 1.51) | OR |
| $1^{-7}$ | 34.5 | 29.0 | 25.5 | 22.9 | 20.8 | 19.2 | 17.7 | 14.0 | $\mu$ |
| | 4.2 | 5.2 | 5.8 | 6.2 | 6.4 | 6.5 | 6.5 | 6.4 | $\sigma$ |
|  | 1.16 (1.01, 1.33) | 1.25 (1.08, 1.44) | 1.28 (1.11, 1.47) | 1.29 (1.12, 1.49) | 1.30 (1.13, 1.50) | 1.31 (1.14, 1.50) | 1.31 (1.14, 1.51) | 1.32 (1.15, 1.51) | OR |
| $1^{-8}$ | 32.4 | 27.0 | 23.6 | 21.1 | 19.1 | 17.5 | 16.1 | 12.6 | $\mu$ |
| | 3.8 | 4.8 | 5.4 | 5.7 | 5.9 | 5.9 | 6.0 | 5.8 | $\sigma$ |
|  | 1.16 (1.01, 1.33) | 1.25 (1.09, 1.44) | 1.28 (1.11, 1.48) | 1.30 (1.13, 1.49) | 1.31 (1.14, 1.50) | 1.31 (1.14, 1.51) | 1.31 (1.14, 1.51) | 1.31 (1.15, 1.51) | OR |

**Table 7.** Hologic C-View odds ratio table: this gives the odd ratios (ORs) for PD_a_ with m = 50 (number of ensemble averages) and n= 2 (detection search window size in pixels) over the range of detection parameters; the top row gives detection parameter 1 and the first column detection parameter 2. There are three metrics cited in each cell: (1) distribution mean (µ, top); (2) distribution standard deviation (σ, middle); and (3) OR with confidence intervals parenthetically. Blue shading indicates possible operating parameters for PDa. The cell with bold fonts and asterisk gives the detection parameters used throughout this report’s examples.

| Detection Parameters | $1^{-1}$ | $1^{-2}$ | $1^{-3}$ | $1^{-4}$ | $1^{-5}$ | $1^{-6}$ | $1^{-7}$ | $1^{-8}$ | |
| --- | --- | --- | --- | --- | --- | --- | --- | --- | --- |
| $1^{-1}$ | 44.7<br>5.8<br>1.36 (1.14, 1.63) | 34.5<br>6.8<br>1.38 (1.15, 1.67) | 28.3<br>6.9<br>1.38 (1.15, 1.67) | **24.0\***<br>**6.7**<br>**1.38 (1.14, 1.67)** | 20.7<br>6.8<br>1.37 (1.13, 1.67) | 18.0<br>6.0<br>1.37 (1.13, 1.66) | 15.9<br>5.6<br>1.36 (1.12, 1.65) | 8.1<br>3.7<br>1.33 (1.10, 1.62) | $\mu$<br>$\sigma$<br>OR |
| $1^{-2}$ | 40.6<br>5.5<br>1.39 (1.15, 1.67) | 31.0<br>6.2<br>1.39 (1.15, 1.68) | 25.2<br>6.2<br>1.38 (1.14, 1.67) | 21.2<br>6.0<br>1.37 (1.13, 1.66) | 18.2<br>5.7<br>1.36 (1.12, 1.65) | 15.8<br>5.3<br>1.35 (1.11, 1.64) | 13.7<br>4.9<br>1.34 (1.11, 1.63) | 6.8<br>3.1<br>1.30 (1.08, 1.58) | $\mu$<br>$\sigma$<br>OR |
| $1^{-3}$ | 38.0<br>5.2<br>1.39 (1.16, 1.68) | 28.8<br>5.8<br>1.39 (1.15, 1.68) | 23.4<br>5.8<br>1.38 (1.13, 1.67) | 19.6<br>5.5<br>1.36 (1.12, 1.65) | 16.7<br>5.2<br>1.35 (1.11, 1.64) | 14.4<br>4.8<br>1.34 (1.10, 1.62) | 12.5<br>4.4<br>1.33 (1.09, 1.61) | 6.1<br>2.7<br>1.28 (1.06, 1.55) | $\mu$<br>$\sigma$<br>OR |
| $1^{-4}$ | 36.2<br>5.0<br>1.40 (1.16, 1.69) | 27.3<br>5.5<br>1.39 (1.15, 1.68) | 22.0<br>5.4<br>1.37 (1.13, 1.66) | 18.4<br>5.1<br>1.36 (1.12, 1.64) | 15.6<br>4.8<br>1.34 (1.10, 1.62) | 13.5<br>4.4<br>1.33 (1.09, 1.61) | 11.6<br>4.1<br>1.31 (1.09, 1.59) | 5.6<br>2.4<br>1.27 (1.05, 1.53) | $\mu$<br>$\sigma$<br>OR |
| $1^{-5}$ | 34.8<br>4.8<br>1.40 (1.16, 1.69) | 26.1<br>5.2<br>1.39 (1.15, 1.68) | 21.0<br>5.1<br>1.37 (1.13, 1.66) | 17.5<br>4.8<br>1.35 (1.11, 1.64) | 14.8<br>4.5<br>1.33 (1.10, 1.62) | 12.7<br>4.1<br>1.32 (1.09, 1.60) | 11.0<br>3.8<br>1.31 (1.08, 1.58) | 5.2<br>2.2<br>1.26 (1.05, 1.52) | $\mu$<br>$\sigma$<br>OR |
| $1^{-6}$ | 33.7<br>4.6<br>1.40 (1.16, 1.70) | 25.2<br>5.0<br>1.39 (1.14, 1.68) | 20.2<br>4.8<br>1.37 (1.13, 1.66) | 16.8<br>4.5<br>1.35 (1.11, 1.63) | 14.2<br>4.2<br>1.33 (1.10, 1.61) | 12.1<br>3.9<br>1.31 (1.08, 1.59) | 10.4<br>3.5<br>1.30 (1.07, 1.57) | 4.9<br>2.0<br>1.25 (1.04, 1.50) | $\mu$<br>$\sigma$<br>OR |
| $1^{-7}$ | 32.8<br>4.4<br>1.41 (1.16, 1.70) | 24.4<br>4.7<br>1.39 (1.14, 1.69) | 19.5<br>4.6<br>1.37 (1.13, 1.66) | 16.1<br>4.3<br>1.34 (1.11, 1.63) | 13.6<br>4.0<br>1.32 (1.09, 1.60) | 11.6<br>3.6<br>1.31 (1.08, 1.58) | 10.0<br>3.3<br>1.29 (1.07, 1.56) | 4.6<br>1.9<br>1.24 (1.04, 1.49) | $\mu$<br>$\sigma$<br>OR |
| $1^{-8}$ | 29.5<br>3.8<br>1.40 (1.16, 1.69) | 21.7<br>3.9<br>1.39 (1.15, 1.69) | 17.2<br>3.7<br>1.36 (1.12, 1.66) | 14.1<br>3.5<br>1.34 (1.10, 1.62) | 11.8<br>3.2<br>1.32 (1.09, 1.59) | 10.0<br>2.9<br>1.30 (1.07, 1.57) | 8.5<br>2.6<br>1.28 (1.06, 1.54) | 3.7<br>1.4<br>1.22 (1.02, 1.46) | $\mu$<br>$\sigma$<br>OR |

**Table 8.** Standardization results when applying PD_a_ to images used for clinical purposes: each larger row contains multiple findings for a given metric. The top row reports β coefficients (unit shift) from conditional logistic regression with 95% confidence intervals (CIs), for each type of transformation, referenced parenthetically: (1) intrinsic pdf (in); (2) transformation to the C-View pdf (cv); (3) transformation to a standardized normal pdf (sn); and (4) z-score transformation (zt). The middle row report odds ratio (OR) for each transformation in per unit shift with 95% CIs with the same labeling as the top row. The bottom row reports the OR for each transformation in standard deviation shift with 95% CIs with the same labeling; a fifth row was added to include the logarithmic transformation (lg) corresponding with the respective OR table. The number of case-control pairs (n) used for the respective calculations is given in the top row. For the combined column n = 180+241+426=847 derived from the respective number of unique pairs in each format.

| Metrics | GE clinical-FFDM<br>n = 180 | Hologic clinical-FFDM<br>n = 667 | Hologic clinical-FFDM<br>n = 241 | C-View<br>n = 426 | Combined<br>n = 847 |
| --- | --- | --- | --- | --- | --- |
| 1. $\beta$ (in) | 0.0384 (0.0170, 0.0599) | 0.0411 (0.0191, 0.0631) | 0.0657 (0.0293, 0.1021) | 0.0488 (0.0202, 0.0775) | 0.0441 (0.0286, 0.0595) |
| 2. $\beta$ (cv) | 0.0704 (0.0311, 0.1097) | 0.0397 (0.0182, 0.0612) | 0.0576 (0.0237, 0.0914) | 0.0488 (0.0202, 0.0775) | 0.0545 (0.0357, 0.0733) |
| 3. $\beta$ (sn) | 0.4657 (0.2054, 0.7259) | 0.2672 (0.1241, 0.4103) | 0.4271 (0.1905, 0.6637) | 0.3262 (0.1348, 0.5176) | 0.3729 (0.2459, 0.4999) |
| 4. $\beta$ (zt) | 0.5074 (0.2325, 0.7824) | 0.2724 (0.1249, 0.4198) | 0.4298 (0.1880, 0.6716) | 0.3418 (0.1452, 0.5384) | 0.3909 (0.2599, 0.5220) |
| 1. OR (in) | 1.04 (1.02, 1.06) | 1.04 (1.02, 1.07) | 1.07 (1.03, 1.11) | 1.05 (1.02, 1.08) | 1.05 (1.03, 1.06) |
| 2. OR (cv) | 1.07 (1.03, 1.12) | 1.04 (1.02, 1.06) | 1.06 (1.02, 1.10) | 1.05 (1.02, 1.08) | 1.06 (1.04, 1.08) |
| 3. OR (sn) | 1.59 (1.23, 2.07) | 1.31 (1.13, 1.51) | 1.53 (1.21, 1.94) | 1.39 (1.14, 1.68) | 1.45 (1.28, 1.65) |
| 4. OR (zt) | 1.66 (1.26, 2.19) | 1.31 (1.13, 1.52) | 1.54 (1.21, 1.96) | 1.41 (1.16, 1.71) | 1.48 (1.30, 1.69) |
| 1. OR SD (in) | 1.59 (1.23, 2.07) | 1.31 (1.13, 1.51) | 1.48 (1.19, 1.83) | 1.39 (1.14, 1.68) | 1.42 (1.26, 1.62) |
| 2. OR SD (cv) | 1.59 (1.23, 2.06) | 1.30 (1.13, 1.50) | 1.44 (1.16, 1.78) | 1.39 (1.14, 1.68) | 1.44 (1.27, 1.63) |
| 3. OR SD (sn) | 1.59 (1.23, 2.07) | 1.31 (1.13, 1.51) | 1.48 (1.19, 1.83) | 1.39 (1.14, 1.68) | 1.45 (1.28, 1.64) |
| 4. OR SD (zt) | 1.62 (1.25, 2.10) | 1.30 (1.13, 1.50) | 1.47 (1.18, 1.83) | 1.40 (1.15, 1.69) | 1.45 (1.28, 1.65) |
| 5. OR SD (lg) | 1.61 (1.25, 2.10) | 1.29 (1.12, 1.48) | 1.52 (1.22, 1.89) | 1.38 (1.14, 1.67) | 1.44 (1.26, 1.63) |

To round out the main standardization analyses, in this subsection we compare measurements from the raw FFDM images with those from the C-view images cited in Table 9. The C-View findings from Table 8 are repeated in this table for ease of comparison. Regression parameters (βs) determined from the various transformations are cited in the top row, noting (1 and 2) are best compared and (3 and 4) best compared. Here, none of the regression parameters were significantly different from zero for the GE-format, in contrast with βs from the Hologic and C-View formats, which are significantly different from zero. The other βs showed variation within and across formats. The middle row shows the ORs in unit shift. When comparing (1 with 2) and (3 with 4), ORs for GE were not significant. The other ORs were both similar and significant within and across formats, restricted to comparison groups. Odds ratios were similar when comparing 1-4 in the bottom row particularly within formats, and the OR for the GE logarithmic transformation in 5 was significant. Similarities between the respective findings determined with PITs and z-transformations suggest the PITs have strong linear trend, as above.

**Table 9.**
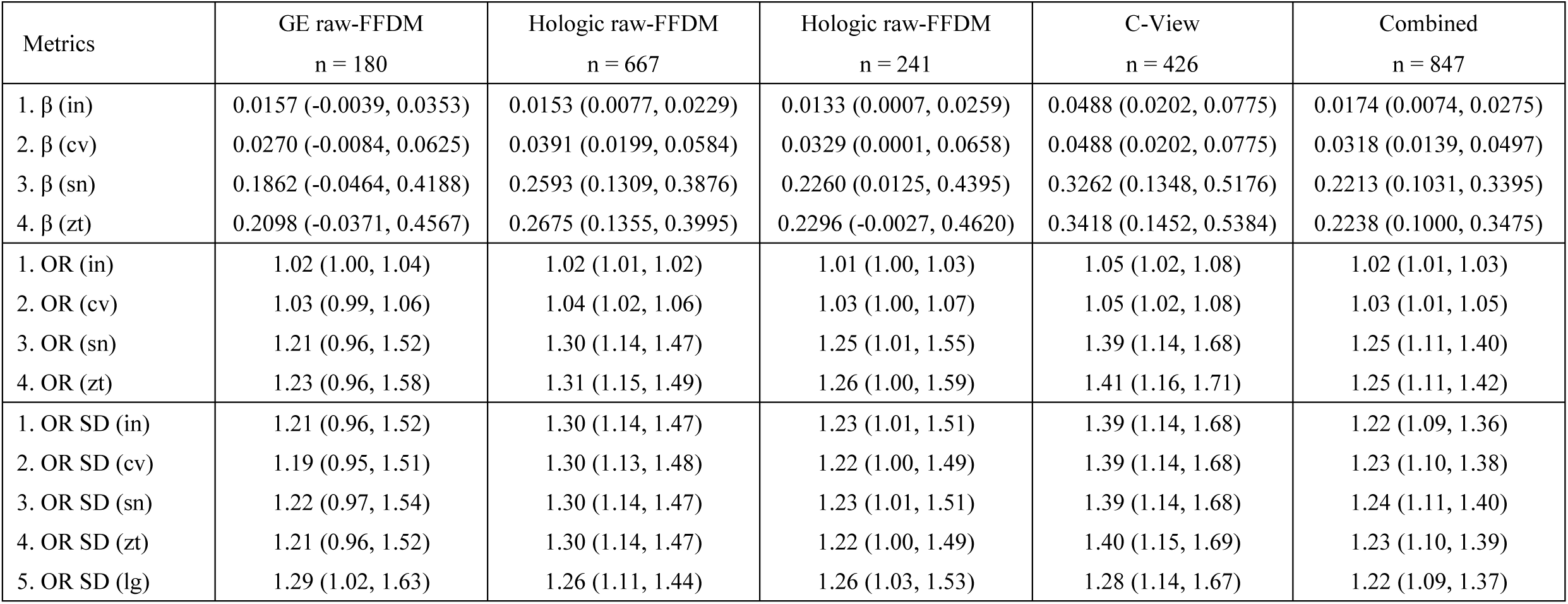
Standardization results when applying PD_a_ to raw FFDM and C-view images: each larger row contains multiple findings for a given metric. The top row reports β coefficients (unit shift) from conditional logistic regression with 95% confidence intervals (CIs), for each type of transformation, referenced parenthetically: (1) intrinsic pdf (in); (2) transformation to the C-View pdf (cv); (3) transformation to a standardized normal pdf (sn); and (4) z-score transformation (zt). The middle row reports the OR for each transformation in per unit shift with 95% CIs with the same labeling. The bottom row reports the OR for each transformation in standard deviation shift with 95% CIs with the same labeling; a fifth row was added to include the logarithmic transformation (lg) corresponding with the respective OR table. The number of case-control pairs (n) used for the respective calculations is given in the top row. For the combined column n = 180+241+426=847, derived from the respective number of unique pairs each format C-view findings are the same as those reported in Table 8 and are repeated here for ease of comparisons.

| Metrics | GE raw-FFDM<br>n = 180 | Hologic raw-FFDM<br>n = 667 | Hologic raw-FFDM<br>n = 241 | C-View<br>n = 426 | Combined<br>n = 847 |
| --- | --- | --- | --- | --- | --- |
| 1. $\beta$ (in) | 0.0157 (-0.0039, 0.0353) | 0.0153 (0.0077, 0.0229) | 0.0133 (0.0007, 0.0259) | 0.0488 (0.0202, 0.0775) | 0.0174 (0.0074, 0.0275) |
| 2. $\beta$ (cv) | 0.0270 (-0.0084, 0.0625) | 0.0391 (0.0199, 0.0584) | 0.0329 (0.0001, 0.0658) | 0.0488 (0.0202, 0.0775) | 0.0318 (0.0139, 0.0497) |
| 3. $\beta$ (sn) | 0.1862 (-0.0464, 0.4188) | 0.2593 (0.1309, 0.3876) | 0.2260 (0.0125, 0.4395) | 0.3262 (0.1348, 0.5176) | 0.2213 (0.1031, 0.3395) |
| 4. $\beta$ (zt) | 0.2098 (-0.0371, 0.4567) | 0.2675 (0.1355, 0.3995) | 0.2296 (-0.0027, 0.4620) | 0.3418 (0.1452, 0.5384) | 0.2238 (0.1000, 0.3475) |
| 1. OR (in) | 1.02 (1.00, 1.04) | 1.02 (1.01, 1.02) | 1.01 (1.00, 1.03) | 1.05 (1.02, 1.08) | 1.02 (1.01, 1.03) |
| 2. OR (cv) | 1.03 (0.99, 1.06) | 1.04 (1.02, 1.06) | 1.03 (1.00, 1.07) | 1.05 (1.02, 1.08) | 1.03 (1.01, 1.05) |
| 3. OR (sn) | 1.21 (0.96, 1.52) | 1.30 (1.14, 1.47) | 1.25 (1.01, 1.55) | 1.39 (1.14, 1.68) | 1.25 (1.11, 1.40) |
| 4. OR (zt) | 1.23 (0.96, 1.58) | 1.31 (1.15, 1.49) | 1.26 (1.00, 1.59) | 1.41 (1.16, 1.71) | 1.25 (1.11, 1.42) |
| 1. OR SD (in) | 1.21 (0.96, 1.52) | 1.30 (1.14, 1.47) | 1.23 (1.01, 1.51) | 1.39 (1.14, 1.68) | 1.22 (1.09, 1.36) |
| 2. OR SD (cv) | 1.19 (0.95, 1.51) | 1.30 (1.13, 1.48) | 1.22 (1.00, 1.49) | 1.39 (1.14, 1.68) | 1.23 (1.10, 1.38) |
| 3. OR SD (sn) | 1.22 (0.97, 1.54) | 1.30 (1.14, 1.47) | 1.23 (1.01, 1.51) | 1.39 (1.14, 1.68) | 1.24 (1.11, 1.40) |
| 4. OR SD (zt) | 1.21 (0.96, 1.52) | 1.30 (1.14, 1.47) | 1.22 (1.00, 1.49) | 1.40 (1.15, 1.69) | 1.23 (1.10, 1.39) |
| 5. OR SD (lg) | 1.29 (1.02, 1.63) | 1.26 (1.11, 1.44) | 1.26 (1.03, 1.53) | 1.28 (1.14, 1.67) | 1.22 (1.09, 1.37) |

In this subsection, we consider an outlier measurement, relative to the historical PBD record (outliers in µ and σ), to investigate its effect on the logistic regression modeling outcomes. We used the GE raw-FFDM images and selected a measurement represented by the appropriate cell in Table 5: specifically, (column 7, row 7) corresponding to the cell with µ= 8.5 and σ = 8.1 (clearly different from the historical expectations, E[µ] ≈ 25 and E[σ] ≈ 17). Different findings were produced relative to the initial results provided in Table 8. These are cited in Table 10, noting these differences: these βs (sn) and (cv) were significant as were the three ORs cited in both the middle (cv, sn, and z) and bottom rows (cv, sn, and lg). Although there were no significant differences between the initial and these OR amplitudes, those cited in Table 10 were consistently larger for all comparisons. This investigation reveals two related characteristics: (1) ORs are a function of the measurement distribution; and (2) the distribution is a function of the algorithm’s operating parameters. The outlier PBD measurement used here produced a valid breast density representation because significant ORs provide an objective truth that it is predictive of risk; this cautiously suggests historical distributions may not be the most relevant. This observation agrees with related work that showed higher density thresholds provided a measure that is more predictive of risk than the corresponding Cumulus application’s thresholds [54]. This analysis also points to the difficulty in establishing a universal breast density measurement. Specifically for PD_a_, these findings show the importance of selecting the appropriate operating parameters and measurement transformation.

**Table 10.** Outlier standardization results for PD_a_ applied to GE raw-FFDM images: we selected outlier operating points relative to the historical findings from Table 5, (column 7, row 7), or the cell with µ= 8.5 and σ = 8.1. Each larger row contains multiple findings for a given metric. The top row reports β coefficients (unit shift) from conditional logistic regression with 95% confidence intervals (CIs), for each type of transformation, referenced parenthetically: (1) intrinsic pdf (in); (2) transformation to the C-View pdf (cv); (3) transformation to a standardized normal pdf (sn); and (4) z-score transformation (zt). The middle row reports the odds ratio (OR) for each transformation in per unit shift with 95% CIs with the same labeling. The bottom row reportw the OR for each transformation in standard deviation shift with 95% CIs using the same labeling; a fifth row was added to include the logarithmic transformation (lg) corresponding with the respective OR table. The number of case-control pairs (n) is given in the top row.

| Metrics | GE raw-FFDM<br>n = 180 |
| --- | --- |
| 1. $\beta$ (in) | 0.0274 (-0.0011, 0.0559) |
| 2. $\beta$ (cv) | 0.0427 (0.0071, 0.0783) |
| 3. $\beta$ (sn) | 0.3126 (0.0642, 0.5610) |
| 4. $\beta$ (zt) | 0.2229 (-0.0093, 0.4551) |
| 1. OR (in) | 1.03 (1.00, 1.06) |
| 2. OR (cv) | 1.04 (1.01, 1.08) |
| 3. OR (sn) | 1.37 (1.07, 1.75) |
| 4. OR (zt) | 1.25 (0.99, 1.58) |
| 1. OR SD (in) | 1.25 (0.99, 1.58) |
| 2. OR SD (cv) | 1.33 (1.05, 1.69) |
| 3. OR SD (sn) | 1.35 (1.06, 1.72) |
| 4. OR SD (zt) | 1.25 (0.99, 1.58) |
| 5. OR SD (lg) | 1.38 (1.08, 1.76) |

Another alternative experiment was used to investigate potential modeling uncertainties that may arise when merging PD_a_ measured from two formats to create a larger dataset. In this analysis, we further investigated the PIT’s effect on the modeling outcome. Measurements from GE clinical-FFDM and C-View images were used represent two similar populations. The two intrinsic distributions for PD_a_ are shown in Figure 3. Figure 4 shows the three relevant PIT approximations: GE clinical-FFDM to normal (left); C-View to normal (middle); and GE-clinical to C-View (right). Figure 5 provides the transformed pdf comparisons. The left pain shows the C-View pdf (black) compared with the corresponding transformed GE clinical-FFDM distribution (red). The right pane shows the normal pdfs for both formats: C-view (black) and clinical-FFDM (red). The intrinsic distribution summaries are cited in the respective OR table (bold fonts and asterisks). For reference purposes, the relevant ORs and CIs in standard deviation shifts are repeated here: (1) GE clinical-FFDM, OR =1.62 (1.25, 2.10); (2) C-View, OR = 1.38 (1.14, 1.67); and (3) transformed GE clinical-FDMM to C-View, OR = 1.59 (1.23, 2.06). Because the number of pairs in the C-View study is considerably larger than the GE study, we randomly sampled 180 CC pairs from the C-View study for this investigation. For the reduced C-View pairs analysis: OR = 1.32 (0.99, 1.76), noting that significance was lost. Combining these two populations (n = 360 pairs) without standardizing their measurements yields OR = 1.80 (1.34, 2.41). Then, combining the populations after transforming the GE clinical-FFDM measures to the C-view distribution produced OR = 1.47 (1.21, 1.77), which appears to be a more feasible finding. Transforming both sets of measures to normal pdfs individually (i.e., an external reference) and combining produced about the same modeling findings yielding OR = 1.47 (1.21, 1.78). Findings from intrinsic C-view measures and those from transforming them to a normal distribution were likely similar because the intrinsic pdf is approximately symmetric. The notable findings are demonstrated by comparing intrinsic ORs from the GE measure and the respective ORs after transforming the measure to either a normal or C-view distribution. The OR from the combined dataset measure without transformation is outside of the upper CI of the combined sample when the PITs were applied. The PITs approximations are shown in Figure 4. As an approximation, PITs appear to have a strong partial linear characteristic over most of the data ranges displayed on the horizontal axes, indicating they are behaving as quasi-linear transformations; although, the curves in the middle and left panes show a nonlinear component as well.

**Figure 3.**
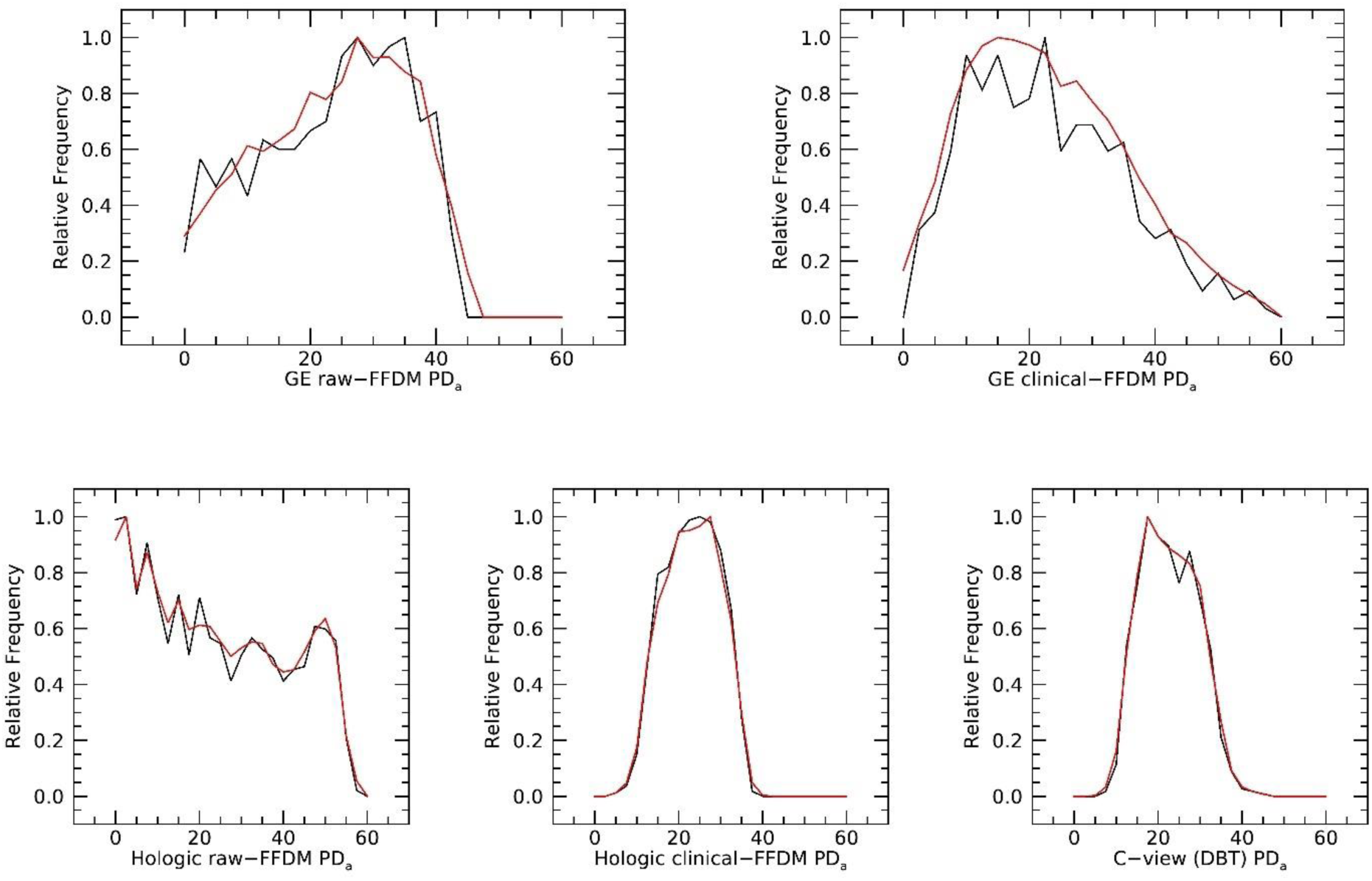
Empirical probability density function (pdf) comparisons with kernel density estimation pdfs: pdfs for PD_a_ distributions are provided for each image format using the detection parameters identified in the odds ratio tables. Solid black lines show the data and red lines pdfs derived from the kernel density estimations, respectively.

**Figure 4.**
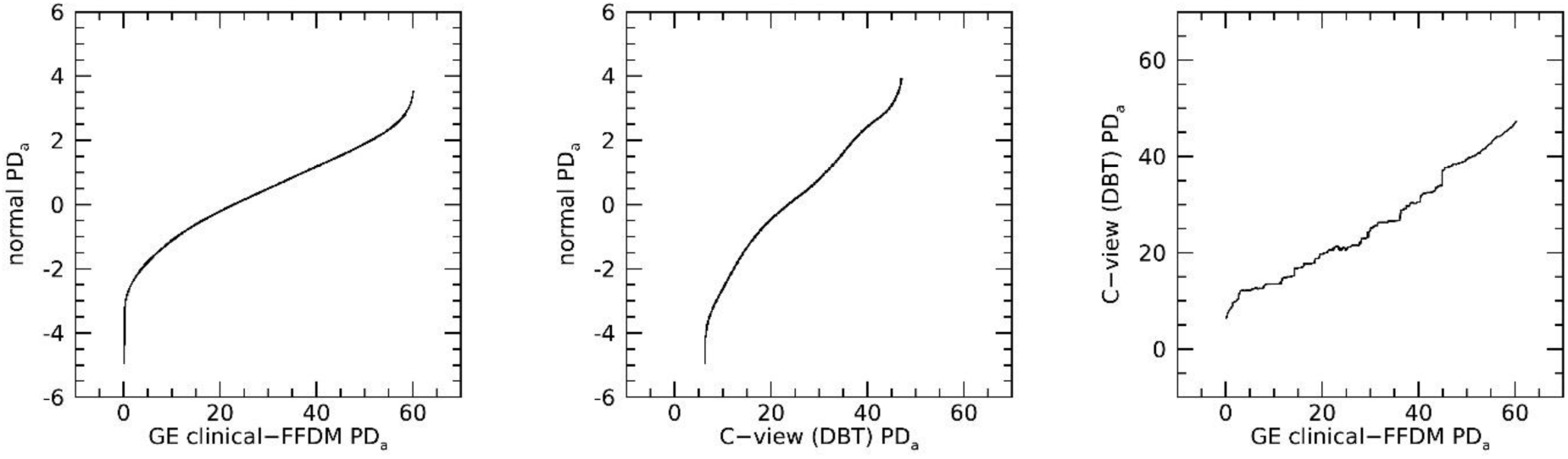
PD_a_ empirical probability density function (pdf) transformation examples for the GE-clinical pdf to C-View pdf: (1) GE intrinsic to normal pdf (left); (2) C-View intrinsic to normal pdf (middle); and GE pdf to C-View pdf (right).

**Figure 5.**
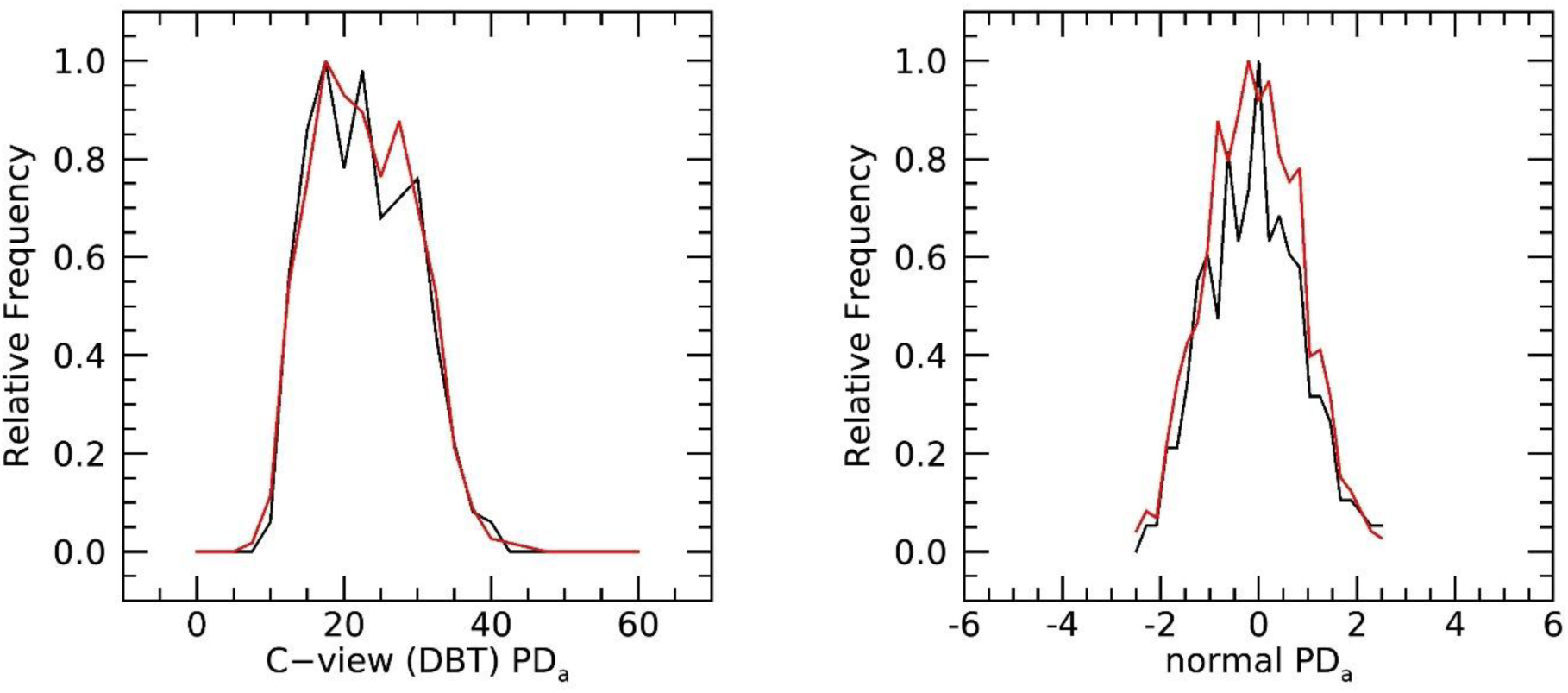
PD_a_ transformation empirical probability density function (pdf) example comparisons: the right pane shows the mapped GE-clinical pdf (red) compared with the C-View pdf (black). The right pane shows both distributions mapped to a standardized normal: GE-clinical pdf (red) and C-view pdf (black).

To summarize the standardization analyses and as a first approximation, the z-score approach may be sufficient to combine breast density measurements across the respective populations. However, reporting negative (less than zero) breast density values in the clinical setting may not be practical. Moreover, converting z-score measurements back to their natural scale can produce inconsistent breast density measurements across populations or image formats. In contrast, the PIT approach overcomes these limitations. The experiment designed specifically to study how to increase n demonstrated that it is possible to combine PBD measurements from different formats (or different studies) using the PIT to form a larger dataset when applying conditional logistic regression modeling. Although further investigations are required, we posit the PIT related findings will apply in the wider modeling context. The outlier analysis also showed that different PBD distributions may be as important as those produced by Cumulus. Although the transformation analysis is a subordinate objective to the main investigation, it is an important complementary component, and we believe such investigations are understudied in the reported literature.

## Discussion and Conclusion

The major innovation of this work follows from synthesizing stochastic processes rather than analyzing mammograms directly. Ensemble averaging over a given participant and the ability to manipulate the exponent p are algorithm advancements permitted by using the synthesized residual image defined in Eq. [2] as the detection input image. Equations [2 and 3] are the SDN transformations that eliminate (makes irrelevant) the requirement for a traditional SDN image as the detection input and can accommodate mammograms that were heavily processed; in previous work, applying a high-pass two-dimensional filter to mammograms was a practical solution to capture an approximation of this residual image. As demonstrated, the SDN relationship can be transformed into a common monotonically increasing quadratic variance structure across the pixel dynamic ranges for different image formats (the same results hold for the raw FFDM images but are not shown). The capability to produce this stochastic quadratic variance structure is essentially image format independent and provides superior variance contrast (p = 1) as reported here. Although only two values for the exponent p were investigated, the results suggest studying values greater than one may provide benefits. The other more subtle algorithmic innovation is the capability to transform a mammogram into statistical state where the chi-square variance ratio test becomes the appropriate detector for dense tissue. This is significant because mammograms inherently exhibit spatially nonstationary statistics.

The opposing effects of ensemble averaging that reduce stochastic variation while enhancing detection performance warrant special comment. In the limit as m → ∞, ensemble averaging collapses the stochastic SDN fluctuation in c_m_(x, y) leaving the deterministic term s(x, y)^2p^. Within this framework, the local variance arises from both noise modulation and local intrinsic structural variation of s(x, y). In the lower extreme (m = 1), excessive variation about the quadratic structure destabilizes detection. Increasing m over a moderate range reduces random fluctuations, which stabilizes the performance. However, increasing m beyond this range suppresses the stochastic SDN component itself, thereby diminishing detection performance. Consequently, the selection of m reflects the balance between reducing excessive variation while preserving the stochastic SDN behavior required for the detection operations. Because the local structure of s(x, y) is format dependent and not affected by ensemble averaging, variability in m was observed.

We adopted a holistic approach to illustrate and select the operating parameters by generating OR tables (tables 3-7) in conjunction with historical measurements. For fixed values of m and p, a given table specifies the problem as a function of the two other detection parameters and gives an overall visual illustration of the number of significant cells. The OR table approach also suggests advantages when applying this detection technique to image data with different formats. In this context, ground truth in the traditional sense may not be necessary; instead, a representative sample from a different image format may be sufficient to determine operating parameters that produce mean values and standard deviations within the ranges reported here.

The simplicity of this density detection formulation is one of its primary strengths compared with more complicated approaches that rely on many free parameters. These attributes may be particularly beneficial for DBT applications. In DBT, the format differences and image appearances are far more varied across manufactures relative to FFDM images due to the respective technical differences in the image acquisitions [55], as well as the introduction of AI enhanced synthetic 2D images (e.g., FDA # K162505: Hologic Intelligent 2D™ Image Reconstruction Software).

There are several qualifications that require mention. Hospital-based populations were studied in the analysis. A small sample from the population of possible image formats was used in this study. However, the evidence reported here demonstrates this approach can adapt, especially for formats other than raw images. Likewise, the number of ensemble averages for a given format was not settled nor was the value for the exponent p. Quantitative methods for determining m for a given image format will be investigated in future studies. In our study, only two values for p were considered, both within the prior constraints. However, letting the exponent p exceed the upper bound (p > 1) will be explored in the future; the variance gain modeling suggests increasing p further may benefit the analysis. We have used these datasets in other breast density analyses so there is always a chance of sampling variation bias and overfitting. However, these concerns are mitigated for several reasons: (1) significant findings were found in all formats investigated over a wide range of operating parameters; (2) the detection process operated on a *tailored* input image, and (3) PD_a_ is limited in the number of free operating parameters.

When considering absolute risk prediction over time, there are unresolved issues that include image measurement and modeling techniques. A few salient points are addressed here. Current clinical-models key in on different measures (with or without density) [8, 56] and do not consider molecular subtypes across breast lesions [57]. There are many automated approaches for estimating breast density. Whether or not a given approach is superior, or preferable for clinical workflow, requires a performance metric for comparison that includes the totality of complexities and requirements for a given approach. For example, a study that compared breast density metrics produced by commercial products with AI that computes PBD [14] illustrates the choice problem when considering the nuances and complexities associated with applying a given technique (e.g., considering free parameters and number of images for training and evaluation). There is limited work in evaluating PBD from DBT data. One AI approach showed ORs were larger from DBT than those estimated from 2D FFDM [58]; this study used the 2D projection images from the DBT acquisition as a starting point to produce the resulting PBD measure (for clarity, 2D projection images are similar to substantially lower dose FFDM images each acquired at a different projection angle that are then used to construct the DBT volume slices). Other researchers found similar findings when investigating PBD estimated from DBT volumes and synthetic 2D images [42]. As the work in this report illustrated, measurement distributions have an impact on ORs making cross-study comparisons difficult, adding to the measure choice problem.

Models used to predict absolute risk may require further investigations to exact how image measurements are included. A given breast density measure is only one component in this process, whereas CNN based absolute risk prediction can incorporate something akin to aggregating multiple image measurements simultaneously (all four views), whether representing a given component in a wider model or as the singular predictor. There is subtlety with CNN based methods that use all four mammographic views. These techniques may be finding early signs of disease if bilateral asymmetry is included in the learning architecture. For example, an AI approach was used to derive breast density and features from benign findings from DBT in a model that predicts short-term absolute risk, with promising findings for screening recommendations [59]. In contrast, the singular breast density measurement approach is most likely insensitive to bilateral differences related to disease. That is, the singular breast density metric is more closely aligned with pure risk prediction when used as a model input rather than risk prediction in combination with early detection. Whether there is an important distinction between risk and risk combined with early detection is not currently evident. It may be for optimized care, different models with varied endpoints could be used in concert for a given patient. Importantly, the appearance of the breast can change over time, more likely shifting to a less mammographically dense state [60] with significant differences between cases and controls [61]. To the best of our knowledge, evaluating the multivariate distribution inclusive of breast density, age and other time changing covariates (e.g., mass or BMI) has not been considered in the absolute risk prediction context to any extent. Current models, accepted for clinical applications, use breast density from a given time point [56]. As mentioned above, to the best of our knowledge, an objective performance comparison metric is lacking that accounts for the complexities required for a given approach to produce a stable prediction over longer-time frames with changing image formats.

Many open-ended measurement/modeling problems will likely be addressed using some form of DBT data. Of special note, including the breast volume slices from DBT to discern an image derived risk factor is a massive data-analytic task, especially when considering four views and age-related distribution changes. We posit side-by-side comparisons of models that use 2D synthetic images versus those the use the respective volume slices are required to document gains (gained precision relative to costs) when considering all of complexities between the two processing tasks. There are multiple models in use now for clinical applications without CNN based AI methods. When considering AI models with the virtually *unlimited* possible variants, developing a standard across institutions and regions fuels an open-ended and required field of inquiry. We are currently developing a large longitudinal retrospective study, where eligible screening participants are selected consecutively. The merits of PD_a_ will be evaluated as a component in multivariate models that predict absolute risk over timeframes and compared with AI models.

## Disclosures

### Author contributions

JH, primary and corresponding author, designed the detection approach, developed code, and directed the experiments. EF is a co-author, assisted in algorithm and investigation designs, performed most of the coding tasks, applied the statistical tests, and developed the graphics in accord with JH MS and KE. MS and KE are co-authors, both of whom provided the required epidemiological expertise for the investigation.

### Internal Review Board

All methods were carried out in accordance with relevant guidelines and regulations. All procedures were approved by the Institutional Review Board (IRB) of the University of South Floridia, Tampa, FL titled, Automated Quantitative Measures of Breast Density: IRB # 104715_CR000002. Mammography data was collected either (1) retrospectively on a waiver for informed consent, or (2) prospectively by gaining informed consent approved by the IRB of the University of South Florida, Tampa, FL under protocol 104715_CR000002.

### Use of Artificial Intelligence

Artificial intelligence (AI) was not used for writing this manuscript, nor was it used for grammatical manuscript proof reading. AI was used in part to search for relevant research articles that were then inspected by the authors for relevance.

### Data Sharing

Image and clinical data used for this work were made public by JH and MS. Data can be found at this reference [62]. Code for this work is not public. The algorithm can be replicated with the details provided in this manuscript, referenced work, and by contacting JH for details.

### Financial and other related disclosures

The authors have no financial or other conflicts to disclose.

